# Stimulus uncertainty and relative reward rates determine adaptive responding in perceptual decision-making

**DOI:** 10.1101/2024.11.14.623587

**Authors:** Luis de la Cuesta-Ferrer, Christina Koß, Sarah Starosta, Nils Kasties, Daniel Lengersdorf, Frank Jäkel, Maik C. Stüttgen

## Abstract

1

In an ever-changing environment, animals must learn to flexibly select actions based on sensory input and the anticipated positive and negative consequences. This type of adaptive behavior is studied using perceptual decision-making (PDM) tasks that feature block-wise changes in stimulus-response-outcome contingencies. Despite extensive research on PDM, there exists no widely accepted mechanistic model of the decision process that captures trial-by-trial adaptation to contingency changes. To address this gap, we first specified three signal detection theory-based models of adaptive PDM. Next, we identified several scenarios in which these models make diverging predictions. For experimental testing, we subjected rats and pigeons to a two-choice auditory discrimination task comprising a sequence of experimental conditions that differed in their stimulusresponse-outcome contingencies. The contingency manipulations were implemented through the concomitant manipulation of reward probabilities, stimulus presentation probabilities and stimulus discriminability across two stimulus-response categories. We find that both rats and pigeons exhibit condition-specific response biases that increase total reward across the entire range of experimental conditions. However, none of the models were able to fit the choice data across all experimental conditions. Through detailed behavior analysis, we demonstrate that learning is driven by the integration of rewards, but not reward omissions. Moreover, model-based analyses reveal that reward integration is influenced by two additional factors, namely perceptual uncertainty and the alignment of steady-state response ratios to relative (rather than absolute) reward differences between the two choices. A model incorporating these factors accounts well for behavioral data across experimental conditions for both species and connects the arguably most influential framework of perception, signal detection theory, with a learning mechanism operating at the level of single trials which, in the steady state, produces behavior consistent with the generalized matching law from animal learning theory.

**Author summary:** Humans and other animals rely on their senses and experience to categorize objects and pursue their goals. For example, a mushroom hunter uses sight, smell and touch as well as knowledge of the local biota to decide whether to pick a particular mushroom. The consequences of erring may be dire – food intoxication if savoring a poisonous exemplar, or a meager dinner if too many palatable mushrooms are rejected. Also, the hunter’s decision may be influenced by ambient lighting conditions or his estimate of how likely it is to encounter poisonous mushrooms in a certain area. Our work is concerned with the algorithms that animals use to make such decisions and how they adapt when circumstances, such as stimulus discriminability, change. We show that mathematical models that incorporate the animals’ uncertainty about the type of stimulus currently being perceived make very similar choices as the animals. Furthermore, we find that the animals balance their choices by considering relative, rather than absolute, reward expectations, reflecting a long-standing principle of animal learning theory. Together, these features collectively allow animals to obtain nearly as many rewards as theoretically possible.

## 3 Introduction

The field of perceptual decision-making (PDM) is concerned with investigating how animals use sensory signals to select appropriate actions. PDM is studied using experimental tasks wherein subjects are required to map (usually two) well-defined responses to two or more well-controlled stimuli. Over the past decades, a substantial body of research has elucidated the neural substrates underlying perceptual decision processes in various sensory modalities and species (10; 26; 48; 51; 55; 58). In parallel with the research on the neural mechanisms of PDM, efforts have been made to characterize the perceptual decision process at the behavioral level. To that end, the responses of subjects engaged in PDM tasks are subjected to rigorous analysis with the objective to characterize trial-by-trial effects of preceding stimuli, responses and outcomes on subsequent decisions. The upshot of this research is that perceptual decisions are not solely determined by the sensory stimuli but also by a multitude of non-sensory influences. These include the exploration of response-outcome contingencies (52) and trial history, i.e., the sequence of preceding stimuli, choices and, outcomes (7; 8; 9; 19; 23; 24; 27); see (1; 4) for more comprehensive treatment.

Notably, in typical PDM experiments, any influence of trial history is detrimental to performance, in the long run reducing the total number of rewards which are gained. This is because animals are confronted with a randomized sequence of stimuli, thus rendering consecutive trials statistically independent and encouraging subjects to emit responses based exclusively on any given trial’s stimulus. Nonetheless, non-sensory factors continue to exert influence on animals’ choice behavior even long after the initial task has been learned. For example (38) showed that reward history influences perceptual decisions even after months of testing in the very same environment, which however may be adaptive in non-stationary environments (41). Several attempts have been made to construct mechanistic models of the perceptual decision process that take trial history effects into account, most of them based on a signal detection theory framework (7; 15; 17; 22; 31; 45; 65), while more recent approaches employ reinforcement learning algorithms (20; 39; 63). However, to our knowledge, there exists neither a consensus on model architectures nor on the precise learning rules that animals employ (e.g., learning from rewards or errors, or both). Signal detection theory (SDT) provides firm ground for developing such comprehensive accounts, first because this framework is frequently used by, and highly familiar to students of perceptual processes, and second because its basic assumptions have been supported in a great number of studies (5; 18; 25; 64) and the theory is continually being tested and extended (29; 34; 66).

The objective of this study was to investigate adaptive responding and its main determinants in a broad range of experimental conditions by applying and developing a set of detection theory-based learning models. To that end, we subjected rats to a PDM task with several experimental conditions featuring concomitant manipulation of reward probabilities, stimulus presentation probabilities, and discrimination difficulty. Building on previous trial-by-trial models (15; 16; 31; 59; 60), we investigated the learning dynamics of subjects and found that obtained rewards, rather than reward omissions, drove adaptation to the new contingencies. Moreover, our findings indicate that the extent of trial-by-trial behavioral adaptation depends on perceptual uncertainty. Additionally, we demonstrate that the steady-state response bias is independent of the total amount of reward received from both sides and only depends on the relative reward rates. This is a well-established finding from animal learning theory (28; 44) which however is rarely connected to models of perceptual decision-making (but see (12; 11; 35)). Finally, we demonstrate that a detection theory-based reward-integration model incorporating these features is capable of fitting and reproducing the behavioral data observed in a second, similarly structured experiment which was conducted with pigeons as subjects.

## 4 Results

### 4.1 Modeling adaptive perceptual decisions in a detection-theory framework

Perceptual decisions made under uncertainty are commonly understood using the SDT framework (22). For the case of a two-stimulus, two-response discrimination task as in Figure 1a, SDT assumes that repeated presentations of the same stimulus generate random values on a decision axis according to a normal distribution. In a single trial, the subject is thus confronted with a random value drawn from one of the two stimulus distributions (S1 and S2) and has to decide which distribution this value was sampled from. The subject indicates that decision by emitting one of two corresponding responses (R1 or R2, respectively). SDT assumes this decision is taken by comparing the observed value of the decision variable with a fixed decision criterion Although SDT itself is silent as to how specific decision criteria are selected or learned, the framework can be extended by specifying how the criterion changes trial by trial as a function of the preceding sequence of stimuli, responses and outcomes. Here, we initially consider three criterion-setting models embodying three straightforward learning rules (Figure 1b). Reflecting their mechanistic structures, these models are named 1) Integrate Rewards (IR), 2) Integrate Reward Omissions (IRO), and 3) Integrate Rewards & Reward Omissions (IR&RO). The operation of these models is exemplified for a series of 5 trials in Figure 1c. A learning agent operating under the IR model only shifts the criterion on rewarded trials (in Figure 1c, trials 1, 3 and 4) as to make the rewarded response more likely to occur in the subsequent trial. In contrast, an agent operating under the IRO model shifts the criterion only in unrewarded trials as to reduce the likelihood of emitting the unsuccessful response again (in Figure 1c, only in trial 2). And third, an agent operating under the IR&RO model shifts the response criterion in both rewarded and unrewarded trials. The models feature one or two learning rates which control the size of the update steps: *δ* for updating after reward in IR and IR&RO models, and *υ* (upsilon) for updating after reward omissions in IRO and IR&RO. Additionally, all models feature a leaky integration of past criteria whose extent is controlled by the leak term γ (the effect of γ is to pull c towards 0 in each trial, its effect is not shown in Figure 1c for simplicity; see Methods section 7.5 for a more detailed explanation).

**Figure 1:**
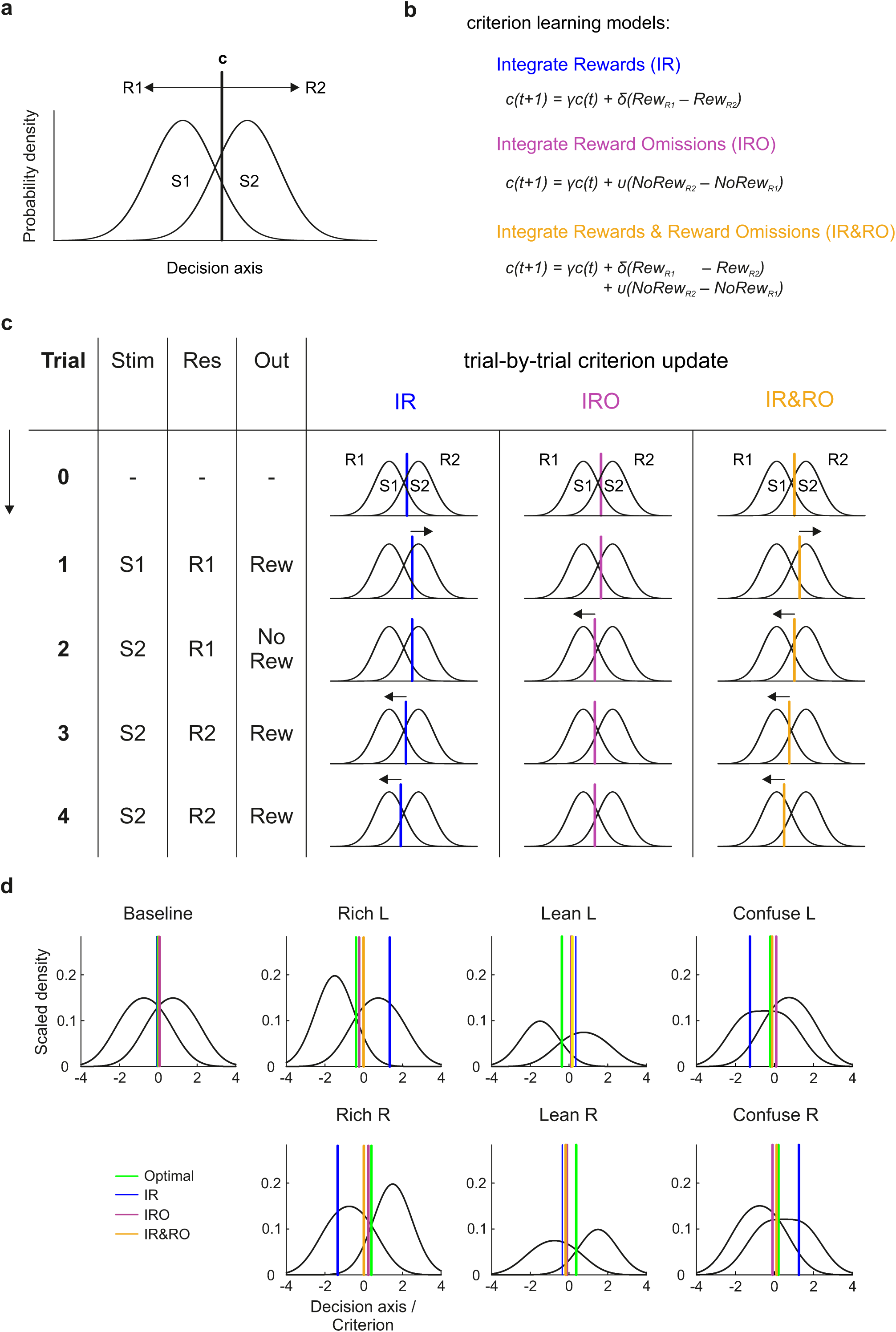
Adaptive decision-making in a detection-theory framework. **a**. Illustration of SDT for a two-stimulus, two-response conditional discrimination task. On any given trial, presentation of a stimulus is tantamount to drawing a random sample x from either of the two distributions (S1 and S2), giving rise to a specific value of a decision variable. The subject makes the decision to emit either response (R1 or R2) by comparing the current value of the decision variable to a criterion c (vertical solid line): R1 if *c < x*, R2 if *c > x*. In classical SDT, the value of the criterion is fixed. **b**. Definitions of three mechanistic models of adaptive criterion setting: 1) Integrate Rewards (IR, blue), 2) Integrate Reward Omissions (IRO, pink), 3) Integrate Rewards and Reward Omissions (IR&RO, yellow). See main text for details. **c**. Exemplification of the criterion updating mechanisms of the three models in a sequence of 4 consecutive trials. Following a stimulus, the subject compares the value of the decision variable generated by stimulus presentation with the current value of criterion c, emits the selected response, receives a reward or not, and then shifts the criterion in accordance with the update rule of each model. The IR model shifts the criterion after each reward, the IRO model after each reward omission, and the IR&RO model after both rewards and reward omissions. **d**. Criterion predictions of the three criterion-setting models and a reward-maximizing account for the steady-state criteria of all experimental conditions. Black lines in each panel represent the ‘decision distributions’, i.e. the category distributions for each of the two categories *z ∈ C*1; *C*2 scaled by presentation and reward probability, i.e. i.e., for category 1, *P* (*x C*1) *P* (*C*1) *P* (*Reward C*1*, R*1) = *P* (*Reward x, R*1) *P* (*x*). Note that the x-value at the intersection of the two decision distributions equals the optimal (reward-maximizing) criterion (see Methods, section 7.5). How these functions are generated specifically in each condition is described in the legend to Figure (S1). (The parameters used for this representation are γ = 0.99*, δ* = *υ* = 0.04).

### 4.2 Experimental conditions to assess the relevance of rewards and reward omissions for adaptive criterion-setting

The three models not only specify exact learning rules for trial-by-trial updating of the decision criterion, but also allow to derive equilibrium predictions for the steady state given a set of parameters (stimulus means, γ, *δ*, and *υ*; see Methods section 7.5 for details). We accordingly designed a set of experimental conditions in which the three criterion-setting learning models predict different steady-state criterion locations. Additionally, we calculated the reward maximizing (henceforth, optimal) criterion location for each condition as a benchmark. Figure 1d presents a general overview of the experimental conditions, Figure S1 provides a more detailed illustration. Generally speaking, there were three sets of conditions dubbed “Rich”, “Lean”, and “Confuse”. Each of the three experimental conditions came in two varieties, L and R. This nomenclature (left, L, and right, R) follows from the criterion shift that would be expected in each of these conditions from an optimal account. L and R varieties of each condition were constructed by mirroring the stimulus locations and their associated reward and presentation probabilities at the baseline category boundary (see Figure S1 for illustration). Crucially, conditions Rich and Lean are scaled versions of each other in terms of the relative amounts of reward to be expected from the two categories, whereas the Confuse condition features one stimulus that is located on the “wrong” side of the category boundary. Additionally, animals underwent repeated testing in a baseline condition which serves as a neutral reference point but does not allow differentiation between the models. For a more in-depth explanation of the specific design of each of the conditions, see Methods section 7.4 and Figure S1. In each condition, rats (N=4) were confronted with three or four auditory stimuli (chords consisting of 11 pure tones differing in frequency content) which were assigned to either of two mutually exclusive categories. The two categories were labeled as ‘low-frequency’ (Category 1, C1) and ‘high-frequency’ (Category 2, C2). Poking into the right side port (Response (R) 1) was considered correct following presentation of a Category 1 stimulus, and accordingly for Category 2 stimuli. However, stimulus presentation probabilities, reward probabilities, as well as the assignment of individual stimuli to the two categories were specific for each experimental condition. Each experimental condition was typically maintained for 10 sessions; and the sequence of conditions was balanced across subjects, with the exception at that the L and R versions of each condition were executed in succession.

### 4.3 Experimental conditions elicit consistent response biases

We subjected rats to a single-interval forced choice (SIFC) auditory discrimination task with multiple stimuli. Rats self-initiated trials by poking into a central response port and had to remain there for 400 ms to trigger presentation of a 100-ms auditory stimulus. Shortly after stimulus offset (25 ms), rats were allowed to withdraw from the center port and indicate their response by entering either of two lateral choice ports (R1 and R2, see Figure 2ab for outline). Premature withdrawal from the center port aborted the trial; aborted trials were not considered for analysis (see Methods section 7.3 for details). Figure 2c shows the fraction of R2 responses (P(R2)) and rewarded trials (P(Rew)) per session for each of the four rats which participated in this experiment. As per design, the Lean conditions (Lean L & Lean R) yielded roughly half as many rewards per session as the Rich conditions (Rich L & Rich R) (Figures 2c and S4). Hit rates and false alarms (calculated for S1 and S5 respectively) changed only slightly over many months of testing (Figure 2d). We first compared the steadystate response bias across experimental conditions. Since stimulus probabilities changed across conditions, P(R2) as shown in Figure 2c is not a suitable measure to perform this comparison, as it confounds response bias and stimulus presentation probabilities. Therefore, we used linear regression to build one-criterion-per-session (OCPS) models which describe performance as resulting from a session-specific criterion and three to five different stimulus distributions which were fixed across all sessions (see Methods section 7.5 and (59; 60). We defined the steady-state criterion as the mean criterion over the last three sessions of each condition for each animal relative to the criterion location in the baseline condition. Per definition, the value of criterion is independent of stimulus presentation probabilities and thus serves as a pure index of response bias (25). As Figure 2e shows, rats consistently shifted their criteria towards negative values (implying a preference for rightward choices, i.e., R1) in conditions Rich L and Lean L whereas towards more positive values in conditions Rich R and Lean R (implying a preference for leftward choices, i.e., R2). Qualitatively, the shift was of comparable magnitude in the rich and lean conditions, suggesting that the different reward densities experienced by animals in these conditions (Figure S4) had no influence on criterion placement in the steady state. In other words, criterion setting was governed by relative rather than absolute reward differences. In the Confuse conditions, animals unexpectedly shifted their criteria towards more positive values not only in the L but also the R variety relative to baseline.

**Figure 2:**
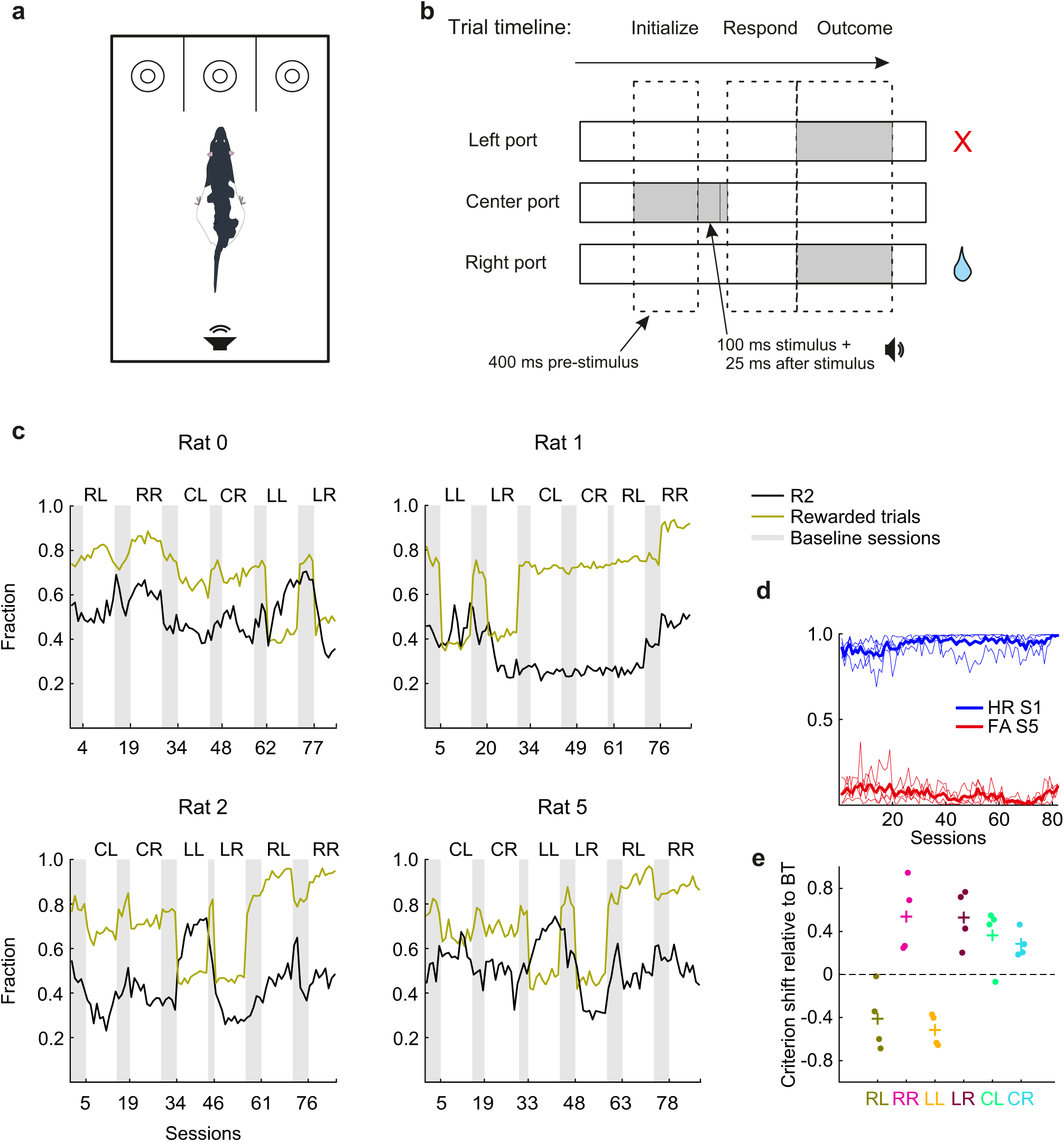
Auditory single-interval forced-choice (SIFC) task and behavioral results. **a.** Schematic drawing of the operant chamber with three conical nose ports. **b.** Schematic outline of the task epochs and possible outcomes. Gray shading indicates continuous poking into a certain nose port. **c.** Response bias (fraction R2, i.e., leftward responses, black) and reward density (fraction of rewarded trials, green) across all experimental conditions for all four subjects. Each panel shows results for one subject, data points represent individual sessions. Conditions are denoted by their respective initials: CL & CR for Confuse Left and Confuse Right; LL & LR for Lean Left and Lean Right, and RL & RR for Rich Left and Rich Right. The gray shaded areas highlight baseline sessions.**d.** Development of hit rate (HR, blue) for stimulus (S) 1 and false alarm rate (FA, red) for stimulus 5 over the course of behavioral testing. Each individual line represents data from a single subject, thick lines represent the means over subjects.**e.** Steady-state criteria observed in the experimental conditions relative to criteria observed in the initial baseline sessions. Points represent the mean steady-state criterion values from the last 3 sessions of each condition for each animal, crosses represent means over the different animals. Observed session-by-session criteria were calculated using the one-criterion-per-session model, see Methods section 7.5 for details.

### 4.4 Rats maximize reinforcement in the steady state

We next asked which of the criterion-setting accounts (IR, IRO, IR&RO, and optimal) best predicts the condition-wise steady-state criteria. To that end, we first fitted rats’ response data with each of the trial-by-trial criterion-learning models and then used the fitted parameters to generate predictions as to where rats’ criteria would converge for each model, given a specific experimental condition and the individually fitted parameters. We then plotted the predicted against observed steady-state criteria (the latter obtained by means of the OCPS model, as above). The results are shown in Figure 3a. While the IR model predictions and IRO model predictions were only weakly (and in the case of IR, negatively) correlated with the observed steady-state criteria (IR: r=-0.56, *r*^2^ = 0.31, IRO: r=0.20, *r*^2^ = 0.38), both the optimal account (r=0.85, *r*^2^ = 0.72) and the IR&RO (r=0.82, *r*^2^ = 0.68) model provided comparably good predictions. To examine whether criterion shifts were adaptive in terms of reward maximization, we compared the absolute distance of the criterion values in the first and the last three (i.e., steady-state) sessions of each condition, relative to the subject-specific reward-maximizing criterion value (Figure 3b). Indeed, rats generally shifted their criteria towards values closer to the theoretical optimum, thereby increasing the fraction of obtained rewards to 97% of the maximally attainable reinforcements (considering imperfect stimulus discriminability and Baseline biases).

**Figure 3:**
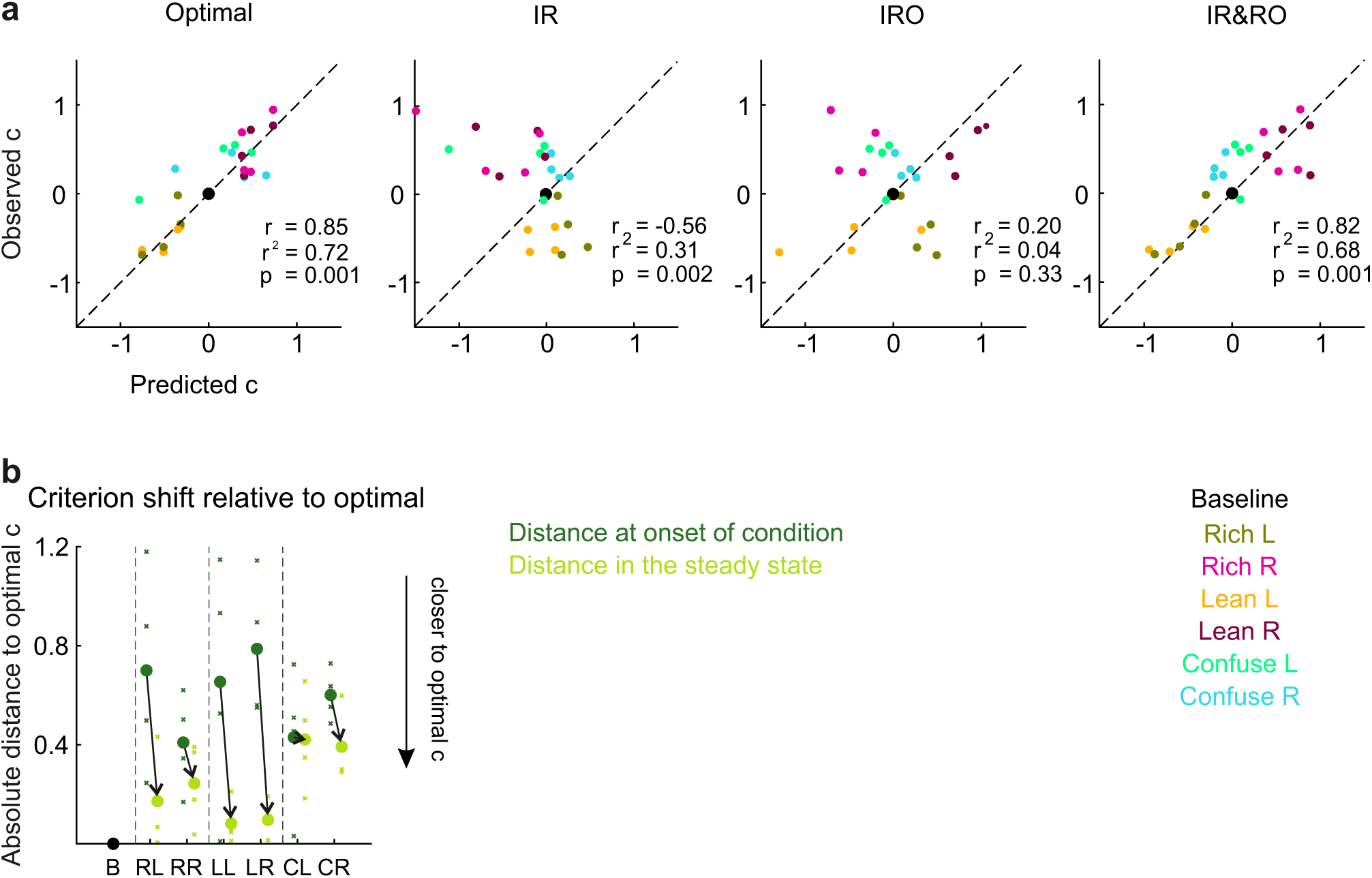
Correlation of predicted and obtained criterion locations in the steady states of the experimental conditions for the rats. **a.** Predicted vs. experimentally observed criterion locations for the Optimal account, IR, IRO, and IR&RO model. Observed criterion locations were computed by means of the OCPS model (see Methods section 7.5) whereas predicted criteria were obtained by fitting each individual model on the whole dataset for each rat, running simulations with the previously fitted parameters (see Methods section 7.6) and averaging over the criteria for all trials within a condition. Individual data points represent a specific pair of predicted and obtained mean criterion for a specific animal in a specific condition. For perfect predictions, data points should fall along the main diagonal (dashed line). Conditions are color-coded. *r*^2^, r and p-values of the correlations are given for each model. **b.** Criterion shift from the onset (average from first three sessions, in dark green) compared to the steady states of the experimental conditions (average from last three sessions, light green), relative to the criterion that would maximize reward in the respective condition. Crosses represent individual animals; points represent means over all 4 animals. All criteria are baseline-normalized prior to plotting.

In summary, all experimental conditions induced response biases whose directions were consistent across subjects. In the Lean and Rich conditions, rats shifted their criteria into opposite directions in L and R versions (moving towards the optimal criterion location), whereas in Confuse conditions rats consistently shifted their criteria towards more positive values in both L and R versions. The latter is likely an undesired artifact of experimental design. The Confuse conditions required extremely tight pre-experimental subject-dependent stimulus selection which post-hoc analysis showed to be suboptimal (see Figure S4b which shows that decision distributions of the Confuse L and Confuse R, arising from suboptimal stimulus means (shown in Figure S5), were almost indistinguishable). We will take up this matter again in Discussion.

Over 10 sessions encompassing several thousand trials, steady-state behavior was tuned towards increasing reward rates close to the statistical optimum. At the same time, response bias neither conformed to the simple IR nor the IRO models, but it aligned with the predictions of the IR&RO model. We next set out to investigate the trial-by-trial performance of these models.

### 4.5 Simple integration of either rewards or reward omissions is insufficient to explain behavior at a trial-by-trial basis

We have so far only looked at steady-state predictions of the learning models. One particular strength of such mechanistic models is their ability to fit choice data on a trial-by-trial basis. We fitted each model to the whole sequence of data for each animal and additionally generated 1000 simulations on a stimulus sequence obtained from a within-condition trial shuffling with the fitted parameters per animal (see Methods section 7.6). Figure 4a shows the results of the trial-by-trial fits of the IR, IRO and IR&RO models to the data for all rats. A model’s performance should be gauged by comparing both fits and simulations to the raw data. A first visual inspection of the IR fits and simulations confirms the results from the steady-state predictions: The IR model generally fails at qualitatively recovering the criterion shifts. As for models IR and IR&RO, although they are better able at fitting the data (as suggested from the steady-state predictions from Figure 3a), the simulations show that these models are also unable to recover the learning trajectories in many of the conditions. Interestingly, inspection of model parameters recovered from the fits revealed that both the IRO and IR&RO model fits featured negative learning rates *υ* for all four animals (Figure 4b). This is at odds as to how these models have been designed to function since it implies that animals shift their criteria as to emit unrewarded responses more rather than less often, and thus compromises the models’ interpretability. The main reason for obtaining the negative learning rates is likely the presence of autocorrelation in the response data: as per design, we induced a response bias, which implies that the probability of the preferred response to occur after any other response is *>* 0.5 in the steady state.

**Figure 4:**
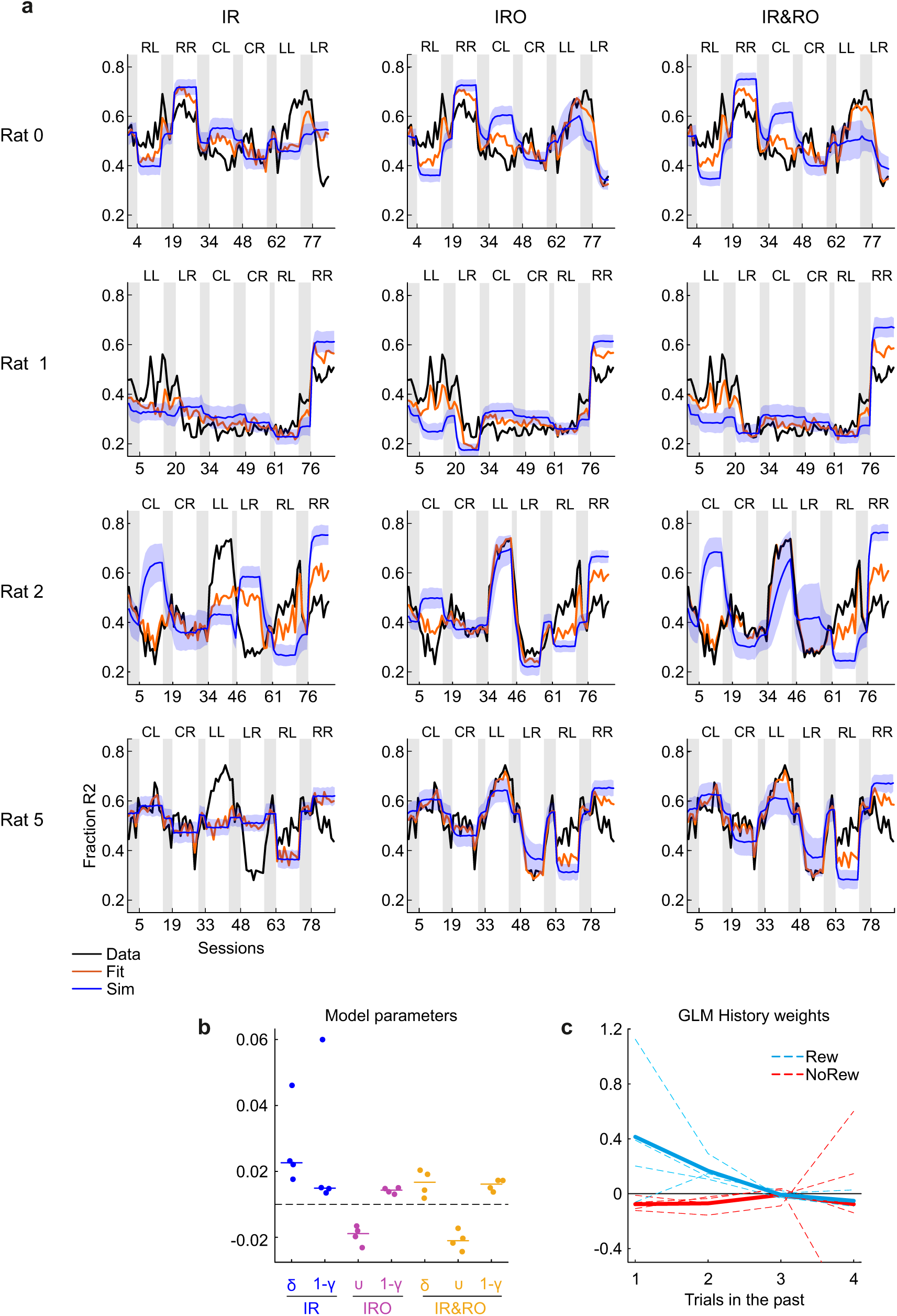
Trial-by-trial fits of the IR, IRO and IR&RO models to the experimental data. **a.** The fraction leftward responses in each session, P(R2), is plotted for each individual animal across the different experimental conditions, similar as in Figure 2c. Orange lines are model fits models and blue lines are averages over 1000 simulations (see Methods section 7.6 for details). Shaded areas represent 1SD. **b.** Distributions of fitted parameters of the different models. Data points represent data from individual subjects. **c.** Regression weights for rewards and reward omissions. The regression weights indicate the influence of both types of outcomes (blue for rewards and red for reward omissions) for trials t-1, t-2, t-3 and t-4. Dashed lines represent individual rats and thick lines means over the 4 subjects. See Methods section 7.5 for details of the GLM fit.

### 4.6 Rewards do affect subsequent behavior while reward omissions have no discernible effect

Due to the inability of all trial-by-trial models to adequately recover response patterns, and the counterintuitive negative *υ* learning parameters returned by the IRO and IR&RO models, we took one step back and analyzed the influence of rewards and reward omissions on subsequent choices from a different perspective. To that end, we regressed the response on trial t as a function of the current stimulus as well as outcomes (rewards and reward omissions) in the preceding trials (t-1, t-2, t-3 and t-4) (27; 40; 46). One would expect rewarded re-sponses to have positive weights (assuming they lead animals towards repeating rewarded responses) and reward omissions to have negative weights (assuming they influence animals towards choosing less often the unrewarded response). This analysis showed that responses were indeed influenced by past rewards whereas the influence of reward omissions was practically nonexistent (on average, regression coefficients for responses following reward omissions where four- or five-fold smaller relative to those of rewards, see Figure 4c). Taken together, the small regression weights, along with the consistently negative *υ* learning rates of the fits returned by the IRO and IR&RO models for all subjects, suggest that reward omissions did not influence the rats’ responses in our task, unlike rewards. We therefore moved on to consider only reward-learning models.

### 4.7 Incorporating stimulus-specific learning rates into the IR model is key to explaining learning trajectories

Confronted with the finding that decisions were indeed influenced by past rewards but hardly by reward omissions with the IR model’s inability to both fit and adequately reproduce learning trajectories in simulations, we were forced to consider additional mechanisms underlying learning from rewards not captured in the original IR model. The learning rate is generally thought to rely on prediction errors, i.e., the size of the difference between expected and obtained outcomes (54); (for example, appetitive discrimination learning proceeds faster with larger rewards than smaller ones (47; 56). In PDM tasks, trials featuring difficult stimuli (i.e., stimuli that lie close to the category boundary) elicit low reward expectations, leading on average to larger prediction errors following rewards, and therefore larger update steps (39; 38). Put differently, learning is maximized after non-predicted outcomes, which can be adaptive in volatile environments (3). Accordingly, to enable the IR model to capture the relation between stimulus uncertainty and update size, we implemented stimulus-dependent learning by fitting a model with one learning rate per stimulus (yielding five learning rates, compared to only one in the original IR model). We coined this extended model Integrate Rewards with Stimulus-specific Learning Rates (IR-SLR). When fitting the datasets with this model, two results became apparent. First, the quality of the fits increased abruptly for all animals (Figure 5a and 5b for an example animal, see Figure S8A and S9B for the others), and forward simulations of this model also qualitatively reproduced the observed behavior in most conditions (Figure 5a, blue shading). Second, as observed by (38; 39) there was a negative correlation between learning rates and discrimination difficulty (the distance of the stimulus means to the category boundary), i.e., rats shifted their criteria by a larger amount following rewards in trials where the discriminative stimulus is close to the category boundary relative to stimuli further away from it (Figure 5c). These results suggest that criterion adjustment is not the same in all rewarded trials, but instead depends on the degree to which a reward is expected based on the discriminative stimulus in that trial.

**Figure 5:**
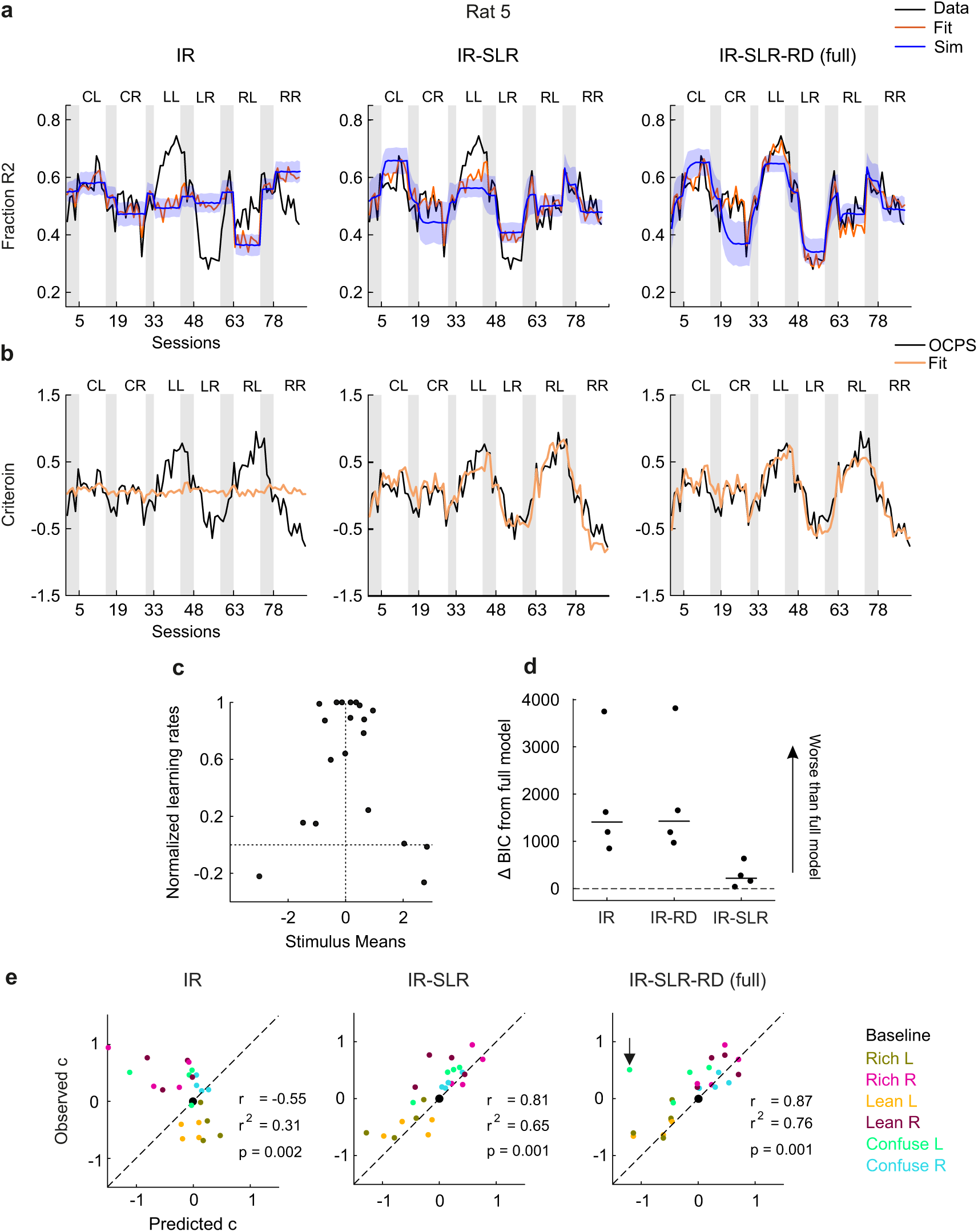
Session-by-session and steady-state performance of the IR, IR-SLR and IR-SLR-RD models. **a.** Fit and simulation results for the two new model versions (IR-SLR and IR-SLR-RD, IR replotted for comparison purposes) for an example animal. Format as in Figure 4a. **b.** Same as in 5a, but plotting session-by-session criteria. **c.** Stimulus-specific learning rates returned by the IR-SLR-RD model as a function of the fitted stimulus means. For comparison purposes, all values were normalized to the overall highest value. Model comparison through the Bayesian Information Criterion (BIC). In this panel, relative values are shown, i.e., the BIC of the IR-SLR-RD (i.e., “full”) model was subtracted from that of the other models, so that positive values are indicative of worse fits than the full model. **e.** Same as in Figure 3a, but comparing steady-state criterion prediction performance of the newly developed models IR-SLR and IR-SLR-RD with the basic IR model. IR-SLR-RD values in plot exclude outlier (highlighted by a black arrow), with outlier values are, r=0.71 and *r*^2^=0.51.

### 4.8 Criterion in the steady state is determined by relative rather than absolute reward differences

While the introduction of stimulus-specific learning rates to the IR model dramatically improves model fits, this model still consistently underestimates the steady-state response bias in the Lean relative to the Rich conditions (Figure 5a, see reduced biases predicted and produced by the models in conditions Lean L and Lean R), which is at odds with our previous observation that animals reached similar steady-state criteria in Lean and Rich conditions (Figure 2e) and which suggested that overall reward density did not seem to affect rats’ steady-state behavior. Importantly, all reward-learning criterion-setting models we discussed so far predict different criteria in the steady state of Rich and Lean conditions (Figure S2). The criterion that the IR model converges to is

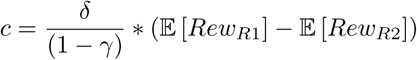

(see derivation in Methods equation 2), so it depends on the absolute difference in reinforcement obtained for R1 and R2. This difference scales with the overall reward density: changing the overall reward density (while keeping the reward ratio between R1 and R2 the same) will also change the absolute difference in reinforcement. For example, when the reward rates for both categories are doubled from one condition to another one, the absolute difference in reinforcement also doubles. Hence, the predicted steady-state criterion for the Rich conditions is higher than the one for the Lean conditions, which have a comparatively lower reward density (Figure S4a). However, as reported above, we found the steady-state criteria to be similar in the Lean and in the Rich condition in our experiment. Accordingly, the IR-SLR requires modification to be consistent with this finding. However, this feature need not be implemented by adding an additional reward density-dependent parameter to control learning rates, but we can instead simply restrict the pull-back uniquely to rewarded trials. In doing so, it can be shown that the steady state to which the model converges to is

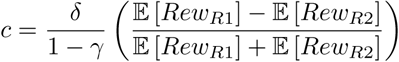

(see Methods equation4). So, the steady is not anymore determined by the absolute but by the relative difference between the rewards obtained from the two categories, which also means that the model becomes insensitive to reward density. Importantly, this minor modification brings the model a step closer to the predictions of the matching law (28), which predicts that the relative response proportion in a two-choice situation is a function of the relative proportion of rewards obtained from the two choices (see Discussion). The new version of the model, dubbed IR-SLR-RD (to make direct reference to the fact that steady-state behavior is controlled by relative reward differences rather than absolute ones), indeed shows clearly improved fits and simulations for the Lean conditions in all rats (Figure 5a and 5b for an example animal, see Figure S8 and S11 for data from other subjects). We quantitatively compared the abilities of all models to fit the whole dataset through the Bayesian Information Criterion (BIC), a measure that takes into account not only the goodness of fit but also the number of free parameters in each model. As expected, the IR-SLR-RD model outperformed all other models (Figure 5d). Lastly, we examined the predictions of the IR-SLR and IR-SLR-RD models for the steady state and found that these predictions were indeed able to both qualitatively and quantitatively capture the steady state of most conditions in all animals (IR-SLR *r*^2^ = 0.65, IR-SLR-RD *r*^2^ = 0.76 (without outlier)), indicating that these modifications together were key to fit rats’ performance particularly when compared with the basic IR model (IR *r*^2^ = 0.31) (Figure 5e).

### 4.9 The IR-SLR-RD model generalizes to pigeons performing a visual PDM task with the same experimental conditions

Quantitative research on adaptive perceptual decision-making is mostly conducted with rats, mice, and humans (see references in Introduction). However, learning per se is of course not restricted to mammals but present in all studied vertebrates (42), and general principles of learning appear to be highly conserved across animals (21; 53). Therefore, to assess the generality of our results, we analyzed data from an additional experiment in which the same battery of experimental conditions was run with four pigeons as subjects (Figure 6a and 3b). The birds performed a structurally similar perceptual choice task which however featured visual (shades of gray differing in luminance) rather than auditory stimuli and a different type of reinforcer, namely access to food instead of water (see Methods section 7.2 for details). All major results from the rat experiments were replicated with the pigeon data. First, neither IR nor IRO models were good predictors of steady-state criteria (Figure 6d), while the combined IR&RO and optimal accounts were better in quantitative terms (*r*^2^) but still far from satisfactory (IR&RO *r*^2^ = 0.15, optimal *r*^2^ = 0.31) Second, the *υ* parameter was consistently negative for all considered models (Figure 6f), suggesting that reward omissions are not a major determinant in adaptive criterion setting. Third, augmenting the IR model with the SLR modification improved the fit in most conditions, although the model was still underestimating steady-state criteria in the Lean conditions (Figure 6e). Fourth, restricting leaky integration to rewarded trials improved the performance in the Lean conditions in the same way as for rats (Figure 6e) and generally provided both excellent fits and forward simulations. Finally, inspection of the BIC values shows that again the IR-SLR-RD model fared best (Figure 6g). Furthermore, this model also exhibited the largest correlation between steady-state predictions and the experimentally obtained criteria (IR-SLR(red) *r*^2^ = 0.52, IR-SLR(red)-RD *r*^2^ = 0.59, Figure 6h).

**Figure 6:**
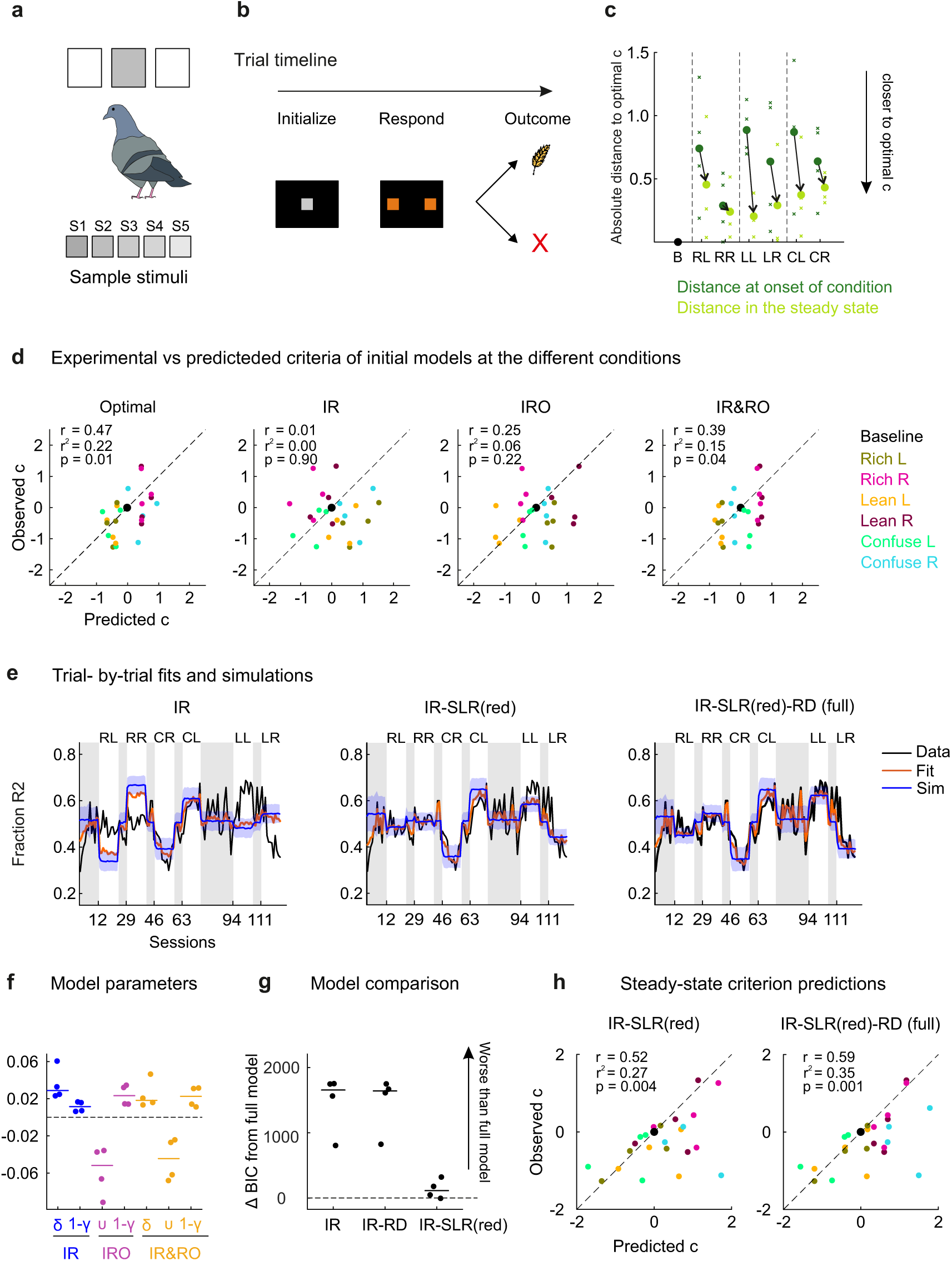
Experimental setup and results from a second dataset from pigeons performing a visual task. **a.** Scheme of the pigeon SIFC luminance discrimination task. **b.** Timeline of an example trial. See Methods for further details. **c.** Criterion shifts (dark green) relative to steady states (light green) for all subjects and experimental conditions. Format as in Figure 3b. **d.** Comparison of the initial steady-state models (IR, IRO, IR&RO and optimal account) in their ability to predict steady-state criteria for all subjects and conditions. **e.** Fits and simulation results of the IR, IR-SLR and IR-SLR-RD models to response data from an example pigeon (subject 897). Format as in Figure 4a. **f.** Model parameters returned by each of the initially considered models. **g.** Model comparison of all the models featuring reward learning. **h.** Steady-state criterion prediction performance of the newly developed models IR-SLR and IR-SLR-RD with the old IR model in the pigeon dataset.

## 5 Discussion

We set out to describe adaptive processes of perceptual decision-making under a broad variety of stimulus-response-outcome manipulations. To that end, we initially considered three different criterion learning models and examined their ability to fit trial-by-trial response data, to generate qualitatively similar data in forward simulations, as well as predict steady-state criterion values. We found that neither of the three models was able to account for the patterns in the data, suggesting that these models are missing essential components. After confirmation that indeed past rewards rather than reward omissions influence choices (above and beyond the discriminative stimuli), we introduced two modifications to the model which have been suggested as key determinants for both trial-by-trial learning and steady-state performance before, but that have so far not been considered in conjunction. These manipulations were first, making learning dependent on stimulus discriminability, and second, making the steady state of the models dependent on relative rather than absolute reward differences across the two categories. These modifications increased the performance of the model with respect to all three aspects of performance (fit, simulations, and predictions of steady-state criteria). Moreover, the importance and generality of these modifications are supported by our finding that they also proffered similar increases in performance in a second dataset collected in pigeons, which were tested with stimuli from a different modality (vision instead of audition) and obtained a different type of reinforcer (food pellets instead of water) but otherwise were subjected to the same experimental manipulations. Interestingly, both features are advantageous in certain scenarios: making learning dependent on uncertainty increases reward rate in volatile environments (3), and matching response ratios to relative reward ratios in the steady state can approximate a reward maximization strategy (67).

### 5.1 Rewards and not reward omissions determine adaptive behavior in our task

We found that fits of all models that incorporated criterion updates after reward omissions (IRO, IR&RO and these same models with the SLR and RD modifications) consistently produced negative learning rates. Under our model architecture and under a lose-switch policy, negative learning rates should not occur as they imply animals shift their criteria as to repeat the last unsuccessful response. We have demonstrated elsewhere that negative parameters are unable to qualitatively reproduce experimental response data in forward simulations (61). Here, we additionally used logistic regression to examine the impact of reward omissions on responding, and confirmed that rewards rather than reward omissions consistently influenced responding in our task. Generally, there are more behavioral states that can lead to an unrewarded trial than to a rewarded one (6; 9; 52). From not paying attention to the sensory evidence during stimulus presentation (9), to fluctuating levels of motivation (6), to exploring uncertain over exploiting known stimulus-response combinations (52), all these are more likely produce unrewarded rather than rewarded outcomes (38). As a consequence, and given the behavioral variability associated with an unrewarded trial, the associations between stimulus-response-outcomes are weaker in trials that end up in reward omissions than those that end up being rewarded. In that sense, our results are in accordance with the literature in that animals display a high degree of response variability after an unrewarded trial which cannot always be attributed to a single learning algorithm (38). Additionally, and specific to some of our experimental conditions, unrewarded trials provide less information than rewards about the currently effective stimulus-response-outcome contingencies. Particularly, in Lean conditions, only 50% of correct responses were rewarded, so animals received definitive feedback on the S-R-O contingency when rewarded but ambiguous feedback when the reward was omitted (because reward implies a correct response, while reward omissions inform only probabilistically on whether the response was correct or not). This particular experimental manipulation may additionally contribute to why reward omissions were not relevant in determining animal performance. Accordingly, future experiments might test the generality of this finding in other designs, e.g., where reward omissions are more informative about the currently effective S-R-O contingency, or where reward omissions are accompanied by aversive stimuli such as foot shocks.

### 5.2 Criterion adjustment depends on stimulus uncertainty and increases expected reward

We found that in both rats and pigeons, reward-learning models could capture and reproduce adaptive choice behavior only after including stimulus-specific learning rates. The fitted learning rates varied systematically as a function of the stimulus means, exhibiting an inverted-U-shaped distribution, with learning rates large for stimuli whose mean was close to the category boundary (featuring high perceptual uncertainty) and small for stimuli whose mean was further away (featuring low perceptual uncertainty; Figure 5c). In perceptual choice tasks such as ours, easily discriminable stimuli are associated with correct responses and subsequent reward delivery more often than more difficult stimuli. According to a reward prediction error framework, rewards following easy stimuli are largely predicted, rewards following more difficult stimuli however are not. (33) demonstrated that rats estimate the perceptual uncertainty of a decision and use it to guide their behavior. In line with this notion, (49) found that in monkeys performing a random-dot motion discrimination task the dopamine neuron activity evoked by discriminative stimuli increases with motion coherence (i.e., decreases with perceptual uncertainty), and the same dopamine neurons were found to signal reward prediction errors. (37) proposed that dopaminergic neurons signal the inverse of perceptual uncertainty, i.e., decision confidence, the degree of belief that a particular stimulus belongs to a given response category. These authors also showed that decision confidence modulates trial-by-trial learning from rewards and continues to do so even after several months of training (38; 39). Importantly, subjects in that latter study performed under constant reinforcement contingencies, so there was no need to adjust performance to varying experimental conditions. Accordingly, in their study any type of adaptation was disadvantageous since consecutive trials were independent. We show here that in the context of an adaptive design in which S-R-O contingencies change dynamically (every one to two weeks constantly over many months of testing), adaptive criterion setting allows the subjects to harvest more rewards, as subjects’ decision criteria move closer to the reward-maximizing criteria in each condition. Indeed, steady-state criterion predictions were highly similar for the optimal and the IR-SLR accounts, in stark contrast to the predictions from the simple IR model in which these predictions were orthogonal to these two accounts (for rats and pigeons respectively, optimal vs IR (r=-0.48, r=-0.19), optimal vs IR-SLR (r=0.69, r=0.58) and Optimal vs IR-SLR-RD (r=0.66, r=0.71)). The fact that this feature of learning is beneficial in the context of an adaptive task and that it is present even when the adaptive component is lacking (as in (38)) suggests that stimulus-specific learning is an integral feature of animal decision-making which is beneficial when S-R-O contingencies are not stationary.

### 5.3 Steady-state biases depend on relative rather than absolute reward differences

Primed by our finding that steady-state criteria in Rich and Lean conditions were of comparable magnitudes, we hypothesized that relative rather than absolute reward differences determine the steady-state biases of subjects. This interpretation is consistent with our previous study in which only reward probabilities were manipulated across conditions (61). We found that the IR model was unable to account for the steady-state criteria while a different model based on earlier work (12) linking signal detection theory to the generalized matching law did successfully predict steady-state criteria (that model is not considered in the present study because it is cannot be applied to data with *>* 2 stimuli). The the matching law posits that choice allocation is a function of relative reinforcement across response options (13) and has been confirmed in a large variety of human and animal decision-making scenarios (28; 43; 44; 61; 62). By modifying the model such that steady-state criteria are independent of reward density by restricting the pull-back to rewarded trials, the model therefore comes closer to the matching law. In the present set of conditions, both the matching law and the IR model with the RD modification predict steady-state response biases which are independent of reward density, consistent with our behavioral results. We acknowledge however that the RD models still differ from the matching law in that their steady-state criterion is directly proportional to the difference in relative rewards, while matching predicts that the criterion is proportional to the difference in their logarithm. We are currently working to develop a trial-by-trial model that is fully congruent with the generalized matching law (35).

### 5.4 Unexplained variance in the Confuse condition

The full model (IR-SLR-RD) provides excellent fits to the choice data for both rats and pigeons. In addition, forward simulations of the model generate similar choice patterns as those obtained experimentally in the baseline, Lean L, Lean R, Rich L and Rich R conditions. However, the model simulations produce discrepant behavior some of the Confuse conditions for some animals (Figure S11, especially Confuse L in rat 2, and Figure S15, pigeons 666 and 902). The main reason for this is an unwanted consequence of the experimental design, namely the high similarity of the two Confuse conditions stimuli 2 and 4) which produced highly similar behavior despite differing S-R-O contingencies. Rats exhibited similar steady-state criteria in the two conditions (Figure 2e). Closer inspection of the decision distributions as well as the objective reward functions in Figure S3 shows that the two Confuse conditions produce highly similar decision distributions on the decision axis, as well as similar optimal decision criteria which, for all four rats, exhibit positive values and therefore lie on the same side of the category boundary. Also, we cannot rule out that the design-inherent confusing S-R-O contingencies in the Confuse conditions, providing ambiguous feedback for stimulus-response combinations due to the similarity of the two central stimuli (S3 and S4 in Confuse L and S2 and S3 in Confuse R, see Figure S1) which however were assigned to different responses, might have led the subjects to “tune out” of the task. Indeed, it has been widely reported that animals leverage different response strategies in PDM tasks associated to varying levels of engagement, or task proficiency (2; 6; 57), and one limitation of our modeling efforts is that this source of variability is neglected.

## 6 Conclusion

In sum, we present a detection-theory model that by design includes the history of stimuli, responses, and outcomes, which together influence upcoming decisions through a single value, the current criterion. Conceptually, the model results bring together and demonstrate the importance of two general features of adaptive perceptual decision-making. These are 1) the inclusion of perceptual uncertainty as a factor which modifies the extent of criterion adjustment and 2) the role of relative rather than absolute differences in reward to determine steady-state response bias. We further report that these two features are particularly beneficial in non-stationary environments, allowing animals to harvest a larger number of rewards. On the light of this finding, we suggest that future research on the mechanisms of PDM as well as its neuronal underpinnings will benefit from incorporating more frequently and more diverse non-stationary contingencies of reinforcement, as are common in natural environments (36).

## 6 Material and Methods

### 6.1 Subjects

#### Rats

Subjects were four male Long Evans rats (Charles River), 6 weeks upon arrival at the institute. Animals were housed in groups of three and lived on a 12-hour reversed day-night cycle (lights off at 8 a.m.). After habituation, they were water-restricted and trained on the behavioral task. Water restriction extended from Sunday to Friday with ad libitum water on weekends. On training days, the animals’ water intake and body weight were measured daily and supplemental water was provided if necessary, with total water intake adjusted to body weight. All subjects were kept and treated in accordance with German guidelines for the care and use of animals in science, and all experimental procedures were approved by local authorities (Landesuntersuchungsamt Rheinland-Pfalz, Germany) and conducted in agreement with directive 2010/63/EU of the European Parliament.

#### Pigeons

Subjects were four unsexed domestic pigeons (Columba livia), obtained from local breeders. Pigeons were housed individually in wire-mesh cages inside a temperature- and humidity-controlled colony room with a light period extending from 8 a.m. to 8 p.m. On weekdays, the birds obtained food in the experimental chambers. On weekends, food was freely available in their home cages. The birds’ body weight was measured daily, and additional food was provided when an animal’s weight dropped below 85% of its free-feeding weight. Water was available ad libitum. All subjects were kept and treated in accordance with German guidelines for the care and use of animals in science, and all experimental procedures were approved by the national ethics review board (Landesamt für Natur, Umwelt und Verbraucherschutz) of the state of North Rhine-Westphalia, Germany, and conducted in agreement with directive 2010/63/EU of the European Parliament.

### 7.2 Apparatus and stimuli

#### Rats

Rats were trained in standard operant chambers (ENV-008, Med Associates) placed inside wooden sound-attenuating cubicles whose interior walls were covered with Styrofoam. Each operant chamber featured three conical nose ports. Nose ports were placed along the side wall and equipped with infrared beams to register snout entries. Also, each nose port featured a small well at its bottom which was connected to a pump for delivery of 30 µl of water. A dim house light was constantly on with the exception of time-out punishments (see below). Sounds were generated in MATLAB (The MathWorks) at a sampling rate of 200 kHz, imported into a custom-written script in Spike2 (Cambridge Electronic Design) which controlled all experimental hardware, then output via a power 1401-3 A/D converter unit (Cambridge Electronic Design) to a conventional stereo amplifier and played by piezoelectric tweeter speakers located at the ceiling of the sound-attenuating cubicle. Following (30), each sound consisted of a chord composed of the sum of 12 logarithmically-spaced pure tones having the same amplitude. Starting from a sound’s center frequency (CF), the 12 pure tones spanned the range CF/1.2 to CF*1.2. Sound duration was 100 ms. For each rat, we selected a set of five chords that lay approximately –1.5, –0.5, 0, +0.5 and +1.5 standard deviations from the category boundary on the decision axis (see Table 1 for each rat’s stimulus frequencies and further below for an explanation of ‘decision axis’). All stimuli were calibrated to 70 dB SPL RMS with a ¼-inch microphone (Microtech Gefell). Feedback sounds (played for time-out punishments and premature withdrawals) were presented from a different speaker which was attached directly to the wall of the operant box.

**Table 1:** Stimulus center frequencies (in Hz) of the chords used for each rat.

| Rat# | Stim 1 | Stim 2 | Stim 3 | Stim 4 | Stim 5 |
| --- | --- | --- | --- | --- | --- |
| 0 | 3249.1 | 5954.5 | 6928.2 | 8061.1 | 14774 |
| 1 | 3249.1 | 6422.9 | 6928.2 | 7473.2 | 14774 |
| 2 | 3249.1 | 5954.5 | 6928.2 | 8061.1 | 14774 |
| 5 | 2792.4 | 5954.5 | 6928.2 | 8061.1 | 17189 |

#### Pigeons

Behavioral testing was carried out in a custom-built operant chamber. The chamber was encased in a sound-attenuating shell; white noise (80 dB) was presented continuously to mask extraneous sounds (Table 2). Visual stimuli were presented on a touch screen (Elo 1515L, Tyco Electronics) mounted to one side of the chamber. A computer-controlled custom-built feeder was located centrally beneath the screen. Upon activation, the feeder provided access to a grain reservoir for 0.5-1.5s (duration was adapted for each animal to ensure adequate food supply depending on body weights and was kept constant throughout the experiment). The chamber was constantly illuminated by two rows of white LEDs positioned beneath the ceiling. Another LED illuminated the food tray during grain delivery. All hardware was controlled by custom-written Matlab code (56).

**Table 2:**
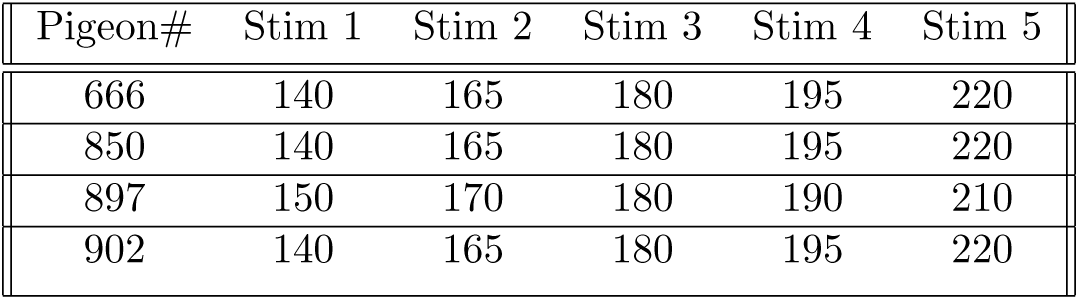
Grayscale values of the visual stimuli used for each pigeon (monitor grayscale values of 140 and 220 correspond to illuminances of 35 and 76 lux, respectively).

| Pigeon# | Stim 1 | Stim 2 | Stim 3 | Stim 4 | Stim 5 |
| --- | --- | --- | --- | --- | --- |
| 666 | 140 | 165 | 180 | 195 | 220 |
| 850 | 140 | 165 | 180 | 195 | 220 |
| 897 | 150 | 170 | 180 | 190 | 210 |
| 902 | 140 | 165 | 180 | 195 | 220 |

### 7.3 Behavioural training and paradigm

#### Rats

Rats performed an auditory discrimination task in which they had to categorize a set of chords as belonging to either a high or low frequency category by emitting a response to the left or right choice port, respectively. During task acquisition, correct responses were always rewarded, and incorrect responses were always punished with a time-out of 4 s during which house lights went off. Once animals reached asymptotic performance (approximately 2 months after beginning of initial training), we mapped the chords to the decision axis. The detection theory concept of the decision axis is illustrated in Figure 1 and explained in the main text. Chords were mapped to the decision axis by converting the fraction of R2 responses for each stimulus to a z-score, which gives the distance of the criterion to the stimulus mean (Figure S3). We then selected, for each subject, five stimuli whose means were located approximately -1.5, 0.5, 0, 0.5 and 1.5 standard deviations from the subjective category boundary (namely, a stimulus that would elicit the same number of R1 as R2). Only these five rat-specific stimuli were used thereafter in the testing phase. Rats performed a single-interval forced choice auditory categorization task in which they self-initiated the trials by continuously poking into the center port for a fixed duration (400 ms), after which the stimulus was played. After stimulus offset, they could immediately withdraw from the center port and emit an operant response (designated as poking into either the left or the right nose port within 4 s; choice phase). Premature withdrawals (i.e., before stimulus offset) led to trial abort, accompanied by a feedback sound and a time-out of 4 s during which the house light was turned off (with the exception of rat 0 for which the time-out was 0 s). Aborted trials were not repeated and not included for analysis. In most experimental conditions, correct responses were consistently rewarded through delivery of 30 µl of water at the selected response port. All non-rewarded responses, correct or incorrect, were punished with a 4 s time-out during which the house light was switched off and a feedback sound was played. Each session lasted 50 minutes and animals completed between 300 and 600 trials. In sessions where animals did not reach a weight dependent-minimum amount of water, they were supplied the remaining volume by the experimenter. During the acquisition phase, the task was designed to provide similar numbers of rewards and reward omissions for each of the categories. During the testing phase, however, rats were subjected to several different experimental conditions, in which they experienced category-wise asymmetrical frequencies of rewards and reward omissions. These asymmetries were brought about by manipulating the mapping between stimuli, responses and outcomes (see Figure S1 further below).

#### Pigeons

Subjects were trained on a single-interval forced choice visual categorization task. Figure 6b provides a sketch of the behavioral paradigm. Key pecks to three distinct rectangular target areas on the touch screen (henceforth, “keys”) were registered as behavioral responses. These three virtual pecking keys were arranged in a horizontal row located about 8 cm above the floor. The central key was positioned in the middle of the monitor, the side keys were placed to the left and the right of the center key. Each trial began with the presentation of an orange-colored rectangle on the center key, accompanied by a 0.5-s pure tone at 1000 Hz. If the animal responded to the stimulus within 3 s (Figure 6b, “Initialize”), one of several discriminative stimuli (shades of gray) replaced the orange rectangle on the center key. Failure to peck at the orange key within 3 s aborted the trial; aborted trials were not repeated. Discriminative stimuli were rectangular uniform gray scale images plotted against a uniform black background. The stimuli only differed in terms of their brightness. Gray scale values ranged from 140 to 220, and were selected according to the discrimination capabilities of individual birds (see Table 2 above). The discriminative stimulus was presented for 1 s and then replaced by an orange rectangle. The birds were required to peck at the orange rectangle at least once within 3 s following discriminative stimulus offset to switch off illumination of the center key and trigger the presentation of two orange rectangles on the side keys. Subjects were required to indicate whether the sample stimulus in the current trial had a gray scale value above or below 180 by pecking at the left or the right choice key, respectively. Correct responses were followed by activation of the feeder for an animal-specific duration. Incorrect responses were followed by a 2-s time-out during which the house light was switched off. The duration of the inter-trial interval (ITI) was 6 s but was extended whenever the birds pecked at the screen within the last second of the ITI until they refrained from pecking for at least 1 s. Testing sessions were conducted on weekdays. Each session consisted of 280 trials and began with three warm-up trials in which the center key was illuminated orange, and a single key peck triggered food presentation. These trials were not analyzed further.

### 7.4 Experimental conditions and model predictions

Animals underwent seven different experimental conditions, termed Baseline (B), Rich Left, Rich Right, Lean Left, Lean Right, Confuse Left and Confuse Right. In the experimental condition **Rich L**, stimulus -1.5 is presented in 50% of trials (C1, consistently associated with R1), while stimulus 0 and stimulus +1.5 (C2) are presented on 25% of trials each and are both associated with R2 (Figure S1). This asymmetry is constructed such that the location of the optimal criterion is to the left of the criterion of the previous symmetric baseline condition. Therefore, an optimal animal should shift its criterion to the left once it enters the new condition (Figure S2). The same is true for an animal that is mostly driven by reward omissions because most omissions with a criterion of zero follow R1 responses, hence making R1 responses less likely by a criterion shift to the left. Importantly, an Integrate-Rewards account predicts that the criterion should be shifted to the right because most of rewards are also obtained from R1 responses and hence R1 responses will become more likely. The upshot of using this stimulus set is that Integrate-Rewards and Integrate-Reward-Omissions learning models make divergent predictions as to the location of the criterion (shifting to positive and negative values, respectively; see S2, predictions for IR and IRO accounts under Rich L conditions). Opposite criterion shifts would be expected if presenting +1.5 in 50% of trials (associated with R2) instead, and presenting -1.5 and 0 in 25% of trials each and reinforcing R1 responses (condition Rich R). See Figure S1 for condition design and S2 for steady state-predictions.

The next condition **Confuse L** follows a stimulus arrangement in which R1 is reinforced ensuing presentations of -1.5 and +0.5 (C1), while R2 is reinforced if occurring subsequent to presentations of 0 and +1.5 (C2; thus, +0.5 is allocated to the “wrong side” of the category boundary). Again, the Integrate-Rewards- and –Reward-Omissions models make divergent predictions about the direction of the animals’ criterion shifts (toward negative values and positive criterion values, respectively). An analogous reasoning applies to condition Confuse R (Figures S1 and S2). Importantly, in both Rich and Confuse conditions, the predictions of each of the two models can be contrasted with an optimization account which predicts criterion shifts in the opposite direction of that of the Integrate-Rewards model in Rich and that of the Integrate-Reward-Omissions model in Confuse. Hence, the two conditions together allow us to diagnose whether animals are more driven by rewards, by reward omissions by both or whether they do something cleverer and can optimize their expected rewards after all.

The last experimental condition considered, **Lean**, represents a replication of Rich in that the ratio of expected rewards from Category 1 vs Category 2 are the same with the exception that subjects are expected to harvest roughly twice as many rewards in Rich compared to Lean versions of the task. That is because in Lean, unlike the previous conditions, reinforcement is probabilistic, and both non-rewarded correct and incorrect decisions trigger time-out punishment. This design allowed us to test adaptation mechanisms underlying different reward-density contingencies. The same experimental conditions were similarly run with both rats and pigeons. Each subject (rats and pigeons) stayed in each condition typically for 10 sessions whereas baseline was run typically for 3 to 5 sessions. In pigeons, the first two sessions of Baseline run between conditions only featured stimulus 1 and 5 (the two easiest discriminable stimuli) whereas the rest of sessions featured stimulus 3 (the stimulus in the category boundary) in 50% of the trials.

### 7.5 Models

#### *Integrate Rewards* model (IR)

The IR model updates the criterion in a stepwise manner only on rewarded trials. Specifically, the criterion in trial t, c(t), is updated according to the following equation:

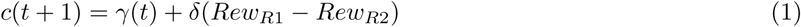

Here, γ ranges from 0 to 1 (usually, 0.9*<* γ *<*1) which represents a leaky integration of past criterion values. With γ=1 (i.e., no leakage), the criterion quickly drifts to infinity (see (50) and (16)). *δ* is a learning rate parameter controlling the size of the criterion adjustment. Following a reward for R1 (*Rew_R_*_1_ = 1, *Rew_R_*_2_ = 0), or for R2 (*Rew_R_*_1_ = 0, *Rew_R_*_2_ = 1), the criterion shifts such that the subject is more likely to choose the same response again in the next trial that led to reinforcement in the current trial. In the steady state, the criterion position of this model depends on the absolute difference in reinforcement obtained from R1 and R2. To understand why, let us derive this criterion position. The model is in an equilibrium when the criterion will not change on average, i.e.: *c*(*t*) = E [*c*(*t* + 1)]. We can compute E [*c*(*t* + 1)] by averaging over all possible outcomes of a trial, weighted with their probability to occur:

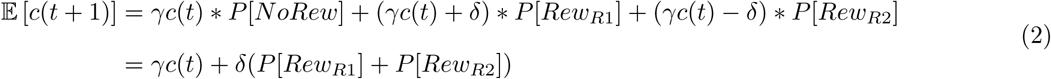

The probability of receiving a reward for a certain response, e.g. *P* [*Rew_R_*_1_], is the same as the expected reinforcement in a trial, E [*Rew_R_*_1_], therefore we can use both terms interchangeably. So, in the equilibrium

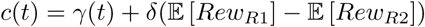

which can be rearranged to

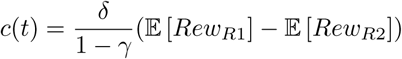

#### *Integrate Rewards* model with convergence to a steady state defined by *Relative Differences* (IR-RD)

This model version follows the same learning design as IR but features no update (including no leak) in the absence of reward. If *Rew_R_*_1_ = 1, or *Rew_R_*_2_ = 1,

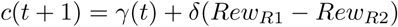

else

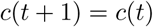

By including this modification, it can be demonstrated that the criterion in the steady state will depend on the difference between relative instead of absolute reward rates. This particular feature makes it more consistent with the matching law, one of the most widely observed equilibriums in decision-making. As before, we can compute the criterion position for the steady state by averaging over all possible outcomes of a trial to get E [*c*(*t* + 1)] and then set *c*(*t*) = E [*c*(*t* + 1)] to determine the equilibrium.

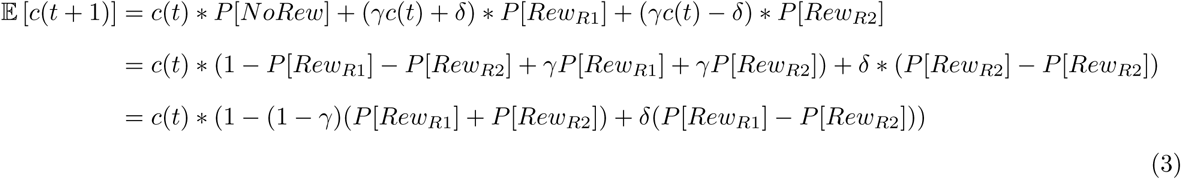

Replacing *P* [*Rew*] with E [*Rew*] and using the equilibrium condition thus gives

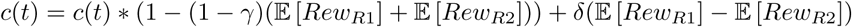

which can be arranged to

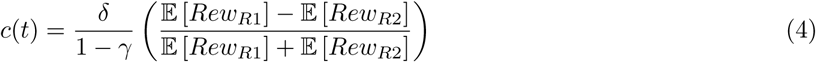

Here, the absolute difference in reinforcement is scaled by the total amount of reinforcement, thereby, removing the dependence on the reward density.

#### *Integrate Rewards* model with *Stimulus-specific Learning Rates* (IR-SLR)

This model is identical to the IR model, with the difference that the learning rate parameter *δ* varies for each individual stimulus. The rationale for using stimulus-specific learning parameters is the hypothesis that animals learn less from events which are more certainly predicted on the basis of sensory evidence. The algorithm is

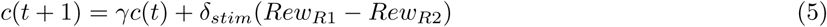

where *δ_stim_* is the stimulus-specific value of *δ* for the stimulus that was presented in trial *t*. Since there were five stimuli over all experiments, there are five values of *δ_stim_*.

#### *Integrate Rewards* model with *Stimulus-specific Learning Rates* and with convergence to a steady state defined by *Relative Differences* (IR-SLR-RD)

As with the IR model the criterion in the IR-SLR model can be designed to converge to a steady state governed by the relative instead of absolute reward differences, in line with the matching law. If *Rew_R_*_1_ = 1, or *Rew_R_*_2_ = 1,

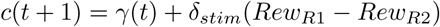

else

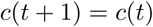

#### *Integrate Rewards* model with a reduced number of *Stimulus-specific Learning Rates* (IR-SLR(red))

This model version is implemented because in pigeons the full IR-SLR model does not provide satisfactory simulations (in 2 out of the 4 animals the simulations go to extreme choice behavior in Confuse L and Confuse R conditions (see Figures S9 and S11). This is because unlike in rats, in pigeons stimulus 2 and stimulus 4 learning rates are fitted only on the base of behavior in Confuse conditions. As shown by the simulations, for some subjects our models are unable to reproduce animals’ behavior in this condition (also see treatment in Discussion). As a result, the learning rates of stimulus 2 and 4 are unreliable. To tackle this, we built a reduced model that we applied to all pigeon datasets, which has only two learning rates, one associated with easy stimuli (1 and 5) and a second learning rate associated with difficult stimuli (2, 3 and 4). This reduced stimulus-specific learning rate modulation is referred to in the manuscript as SLR(red) and does not qualitatively alter model performance although it features less degrees of freedom (see Figure 6g and Figure S7e).

#### *Integrate Reward Omissions* model (IRO)

The IRO model follows the same logic as the IR model, the main difference being that it updates the criterion only on unrewarded trials. For reward omissions following R1, (*NoRew_R_*_1_ = 1, *NoRew_R_*_2_ = 0), and vice versa reward omissions following R2, i.e., *NoRew_R_*_1_ = 0, *NoRew_R_*_2_ = 1. The model renders the unrewarded response less likely to occur in the following trial by shifting the criterion as described by:

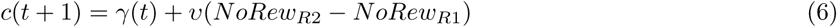

The size of the criterion step is now controlled by the learning parameter *υ*. Negative learning rates imply a tendency of the model to increase the probability to choose the response that leads to a reward omission.

#### *Integrate Rewards & Reward Omissions* model (IR&RO)

The IR&RO model updates the criterion in both trials with rewards and reward omissions according to:

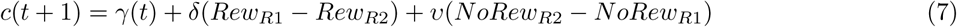

The size of the criterion step after rewards is controlled by *δ* whereas after reward omission it is controlled by *υ*. The IRO and IR&RO models were expanded to encompass stimulus-specific learning and/or converge to a steady state defined by relative differences, but as shown in Figure S6a (rats) and S7e (pigeons), the fits still yielded negative learning rates and were therefore not considered in the main body of the paper.

#### Criterion setting according to an optimal account

To benchmark the animals’ performance, we computed the optimal location of the criterion within the SDT framework as a function of the fitted stimulus distribution means and the experimental reward contingencies for each condition. The optimal criterion maximizes the expected reward

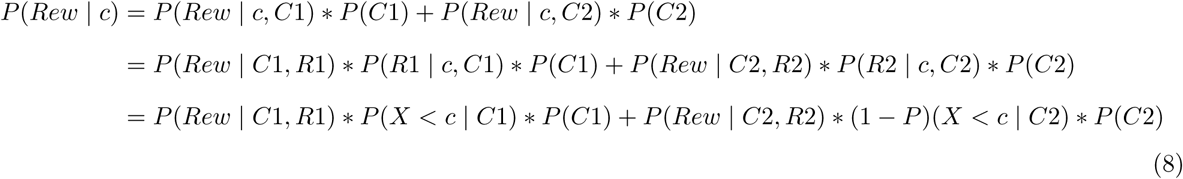

To maximize this, the first derivative with respect to c needs to be zero, which is equivalent to

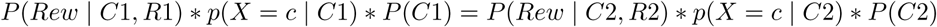

We solved this equation through numerical optimization. Conceptually, it means that the optimal criterion is located at the intersection of the category distributions scaled with the presentation and reward probability (see Figure 1d for a visualization). Note that the scaled category distribution can be obtained by summing the scaled stimulus distributions for all stimuli that belong to that category, i.e., for category 1,

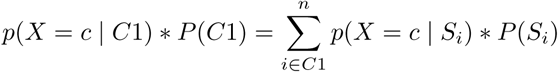

We use the term ‘decision distribution’ as a shorthand for the scaled category distribution because it summarizes all three decision factors (*x*, *P* (*Reinf*), and *P* (*Stim*)) and can be interpreted as the function of observation-dependent action values, scaled by the probability of the respective observation.

#### *One-criterion-per-session* model (OCPS)

Because we manipulated the stimulus presentation probabilities (fraction of trials that belonged to each category) across conditions, the R2 probabilities do not directly reflect the decision criterion but partly follow a different trajectory. We therefore show the behavioral data not only as P(R2) but also as criterion. In contrast to P(R2), the criterion is not directly observable from the behavior. To determine the criterion, we modeled the animals’ stimulus-wise R2 probabilities for each session as a function of stimulus means (one for each stimulus, remaining constant over the course of the experiment) and a session-specific criterion. Using the standard signal detection theory model and assuming a fixed criterion for each session, the probability of choosing R2 in session *j* when stimulus *i* was presented can be expressed as:

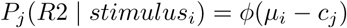

where *ϕ* denotes the cumulative normal distribution, *µ_i_* is the mean of the distribution of stimulus *i* and *c_j_* is the criterion in session *j*. In our one-criterion-per-session model, we thus computed the z-scored probabilities of responding R2 for each stimulus *i* and session *j* (*d_ij_*) by taking the inverse cumulative normal distribution *ϕ*^−1^ of the observed response probabilities.

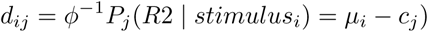

This gives us a model that has one parameter *µ_i_* per stimulus and one parameter *c_j_* per session, representing the criterion for each respective session, which we fitted with linear least-squares regression, using dummy coding for the stimuli and sessions.

#### Logistic regression analysis

We built a logistic regression model (using the glmfit function in Matlab, assuming binomially distributed responses and using the logit link function) to investigate the impact of past rewards and reward omissions on the subsequent choice probabilities in rats. We regressed the influence of previous rewards (*Rew*) and reward omissions (*NoRew*) in trials *t* − 1, *t* − 2, *t* − 3 and *t* − 4 on the response (0 for R1 / 1 for R2) in trial *t*. All the history regressors were built so that the propensity towards R2 was coded as 1 and towards R1 as -1. Specifically, both reward for R1 and no reward for R2 were coded as a -1 in the *Rew* and *NoRew* regressors respectively, whereas reward for R2 and no reward for R1 were both coded as +1. We additionally included dummy-coded regressors for each stimulus *i* (one regressor per stimulus), as well as one regressor for each session *k* to account for slow criterion changes (excluding the first session to avoid collinearity of regressors). Only after completion of the regressor table, we excluded aborted trials from analysis, thereby removing them from the dependent variable, but still having an indirect impact in the history regressors (e.g., if an aborted trial happened at t-1, and although that given trial would not be considered, *Rew_t_*_−1_ and *NoRew_t_*_−1_ regressors would incidentally both equal 0 at row *t*).

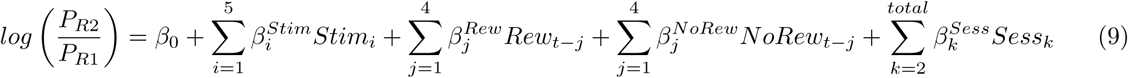

### 7.6 Model fitting and forward simulations

The model fitting was performed as described in (60). To summarize, for a fixed leak parameter (γ), the models can be expressed as a generalized linear model. (14) shows that the likelihood function of these models has a unique maximum which can therefore be found using standard numerical optimization methods. The models were fitted by repeating this procedure for different values of γ and choosing the parameters leading to the overall maximum likelihood. We compared the goodness of fit of the different criterion learning models through calculations and comparison of the Bayesian Information Criterion (BIC) values for each of the respective model fits:

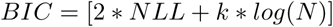

where *k* is the number of free parameters of the model, *N* the number of trials and *NLL* is the negative log likelihood of the observed data given the model and fitted parameters. The advantage of using the BIC for model comparison rather than NLL is that the BIC controls for the number of free parameters, which may differ between models. Models with smaller BIC values are preferred. According to (32), the strength of evidence against the model with the higher BIC value is “positive” for BIC differences of 2-6, “strong” for differences of 6-10, and “very strong” for larger differences.

In each simulation, the models were presented with a newly generated sequence of stimuli and potential rewards (i.e., sequence of trials where a reward will be acquired given a correct response), which was obtained from shuffling all the trials within the same condition. For each trial, the model’s response was sampled according to its predicted probability for each response, and then the criterion was updated according to the outcome following the respective model’s update rule. For each animal and model, we ran 1000 simulations and computed the mean and standard deviation of the fraction of R2 responses per session. Note that in order to run simulations with models featuring SLR modulations in pigeons we had to implement a reduced form with only 2 rather than 5 fitted learning rates (coined IR-SLR(red) model, see Models section above). The results with 2 or 5 learning rates provided comparably similar improvements as depicted by the BICs differences (Figure 6f and S7c), see justification of this approach above in Methods section 7.5).

## 8 Acknowledgements

Preparation of this work was supported by grants from the Deutsche Forschungsgemeinschaft to M.C.S. (project IDs 197059818 and 424828846) and F.J. (project ID 424828846).

The authors thank Alex Hyafil for the assistance with the logistic regression model used to investigate the influence of rewards and reward omissions on subsequent choices.

## 10 Supplementary figures

**Figure S1:**
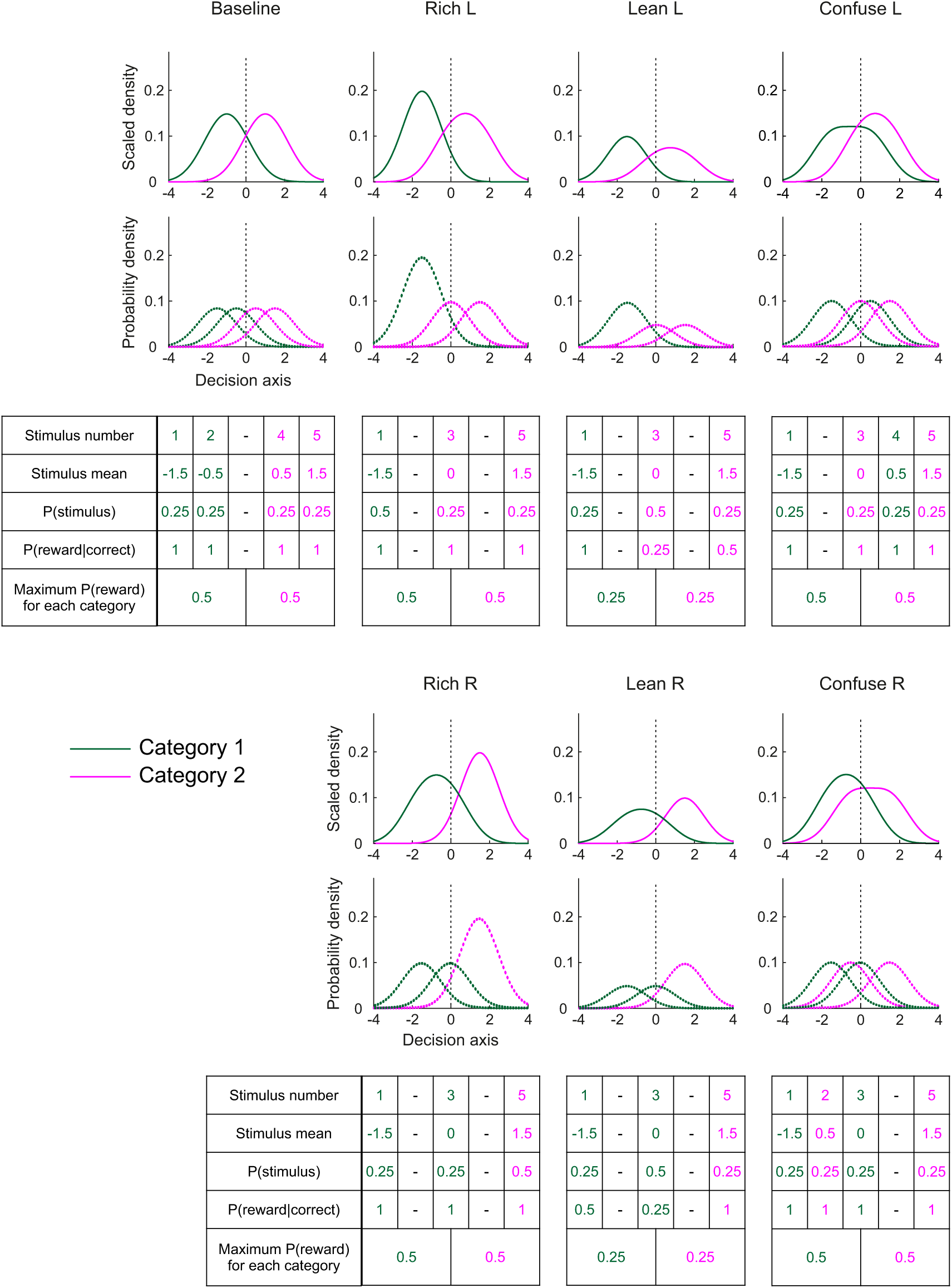
Breakdown of the experimental design of conditions into the specific stimuli, stimulus presentation probabilities and reward probabilities associated with each category. For both figure panels and tables, green lines and font represent category 1 whereas pink represents category 2. For each condition, the dotted lines in the bottom panels represent the stimulus distributions, while the thick lines in the top panels represent the decision distributions for each category (see legend to Figure 1d for explanation). Pigeons Baselines featured slightly different stimulus set (featuring stimulus 3 (P(stim)=0.5) instead of 2 and 4 (P(stim)=0.25+0.25) as in rats).

**Figure S2:**
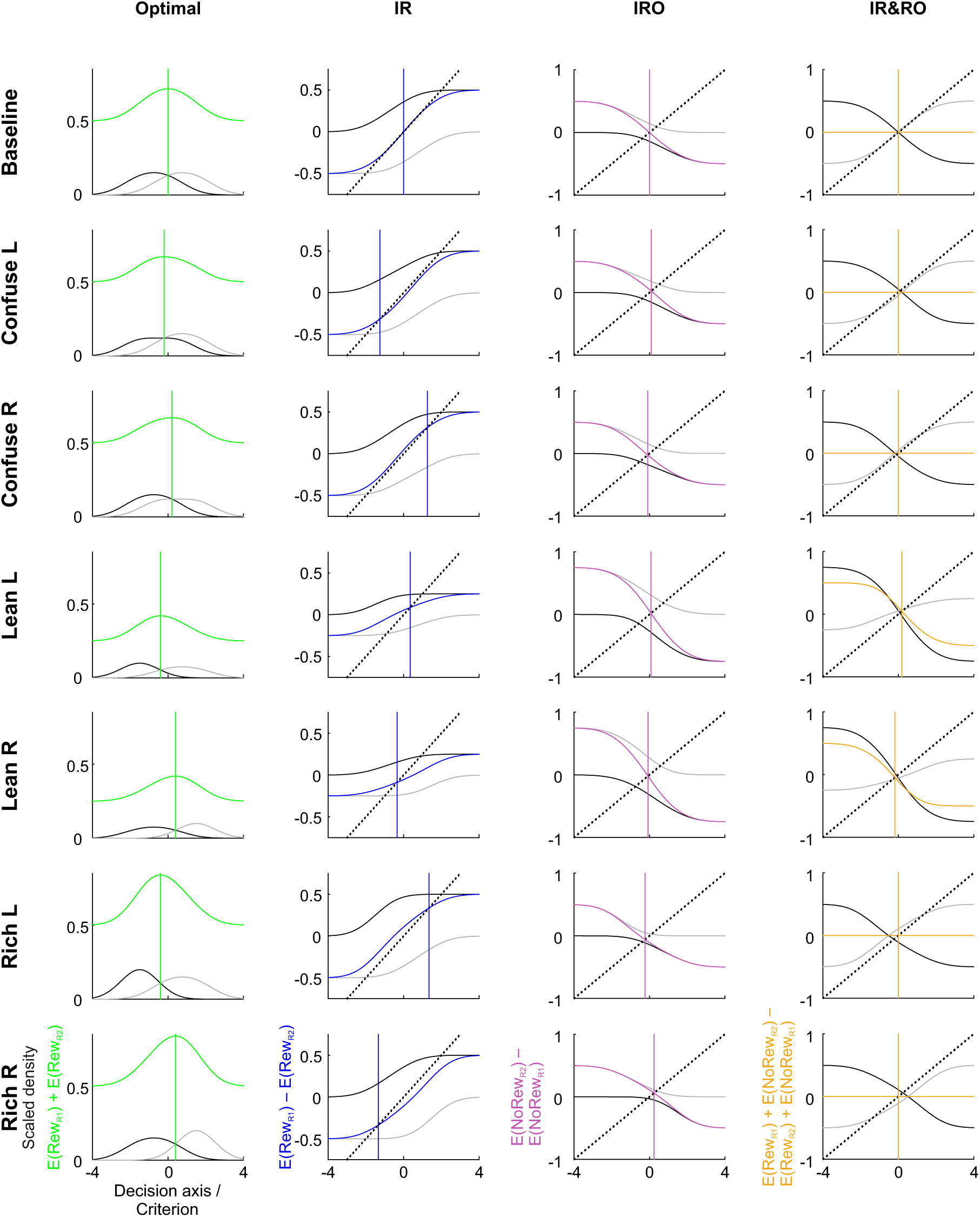
Exemplary predictions of steady-state criterion locations for the three initially considered models for all experimental conditions. Each row shows the predicted criterion locations (vertical colored lines) of the optimal account, IR, IRO and IR&RO accounts for a certain experimental condition. In the first column, the optimal account is shown. In all plots in this column, the green curve is the objective reward function, which represents the total expected probability of reinforcement in a trial dependent on the criterion position. The black and gray lines are the decision distributions for R1- and R2-associated stimulus categories, respectively. Optimal performance is achieved at the maximum of the objective reward function; the corresponding criterion is plotted as a green vertical line. In the other columns, predictions for the IR, IRO and IR&RO models are shown. The colored lines depict the category-wise differences between the expected probabilities for reward (IR), reward omissions (IRO), or both (IR&RO). The black and gray lines depict the same, but conditioned on a trial with R1 and a trial with R2, respectively. Additionally, a dashed black line through zero is plotted, whose slope depends on the leakage term γ and the step size *δ* or *υ*: (1–γ)*/δ* for the IR model, (1–γ)*/υ* for the IRO model, and (1–γ)*/δ* = (1–γ)*/υ* for the IR&RO model. The predicted criterion location for the models is at the intersection of this straight line with the colored line (see Methods section for the derivation). Parameter values used for these predictions:γ = 0.99*, δ* = *υ* = 0.04.

**Figure S3:**
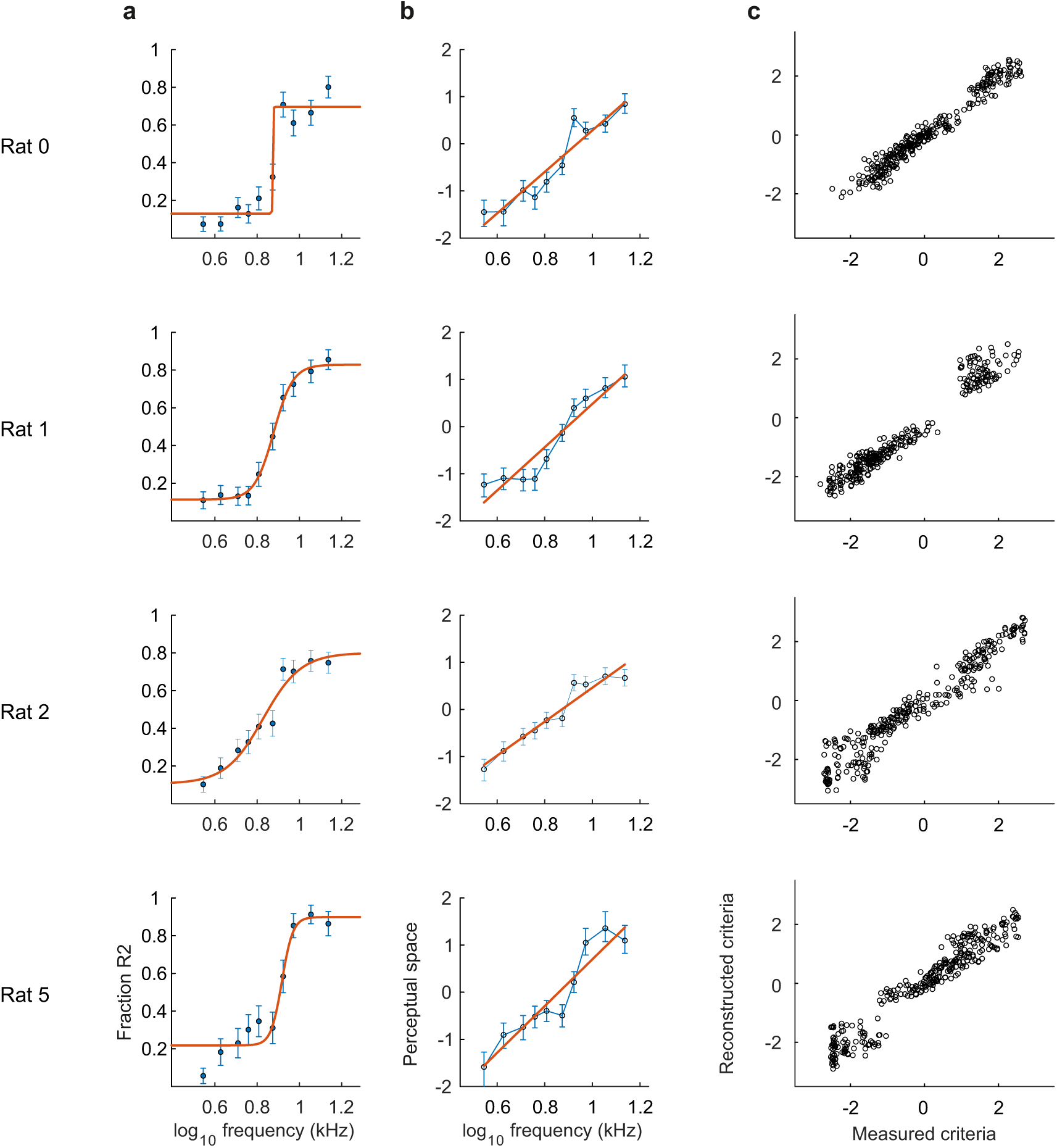
Subject-specific construction of stimulus sets and validation of SDT framework. **a.** Psychometric functions of each rat, fitted with logistic functions and allowing for lapses (orange). Each blue data point represents the fraction of leftward choices for one stimulus. Bars represent standard errors of the means. **b.** Same as a, but after z-scoring P(R2) values to map the stimuli to perceptual space (0 corresponds to the stimulus for which the subject will respond R1 or R2 with equal probability). This mapping was carried out to select a suitable set of stimuli that would match as closely as possible the location of the stimulus means in the perceptual space specified by the experimental design (-1.5, -0.5, 0, 0.5, 1.5). c. Each point represents the reconstructed criteria (elicited by stimulus *i* in session *j*), as calculated from the OCPS model vs the z-scored choice probabilities (elicited by stimulus *i* in session *j*), directly measured from each of the rat subjects.

**Figure S4:**
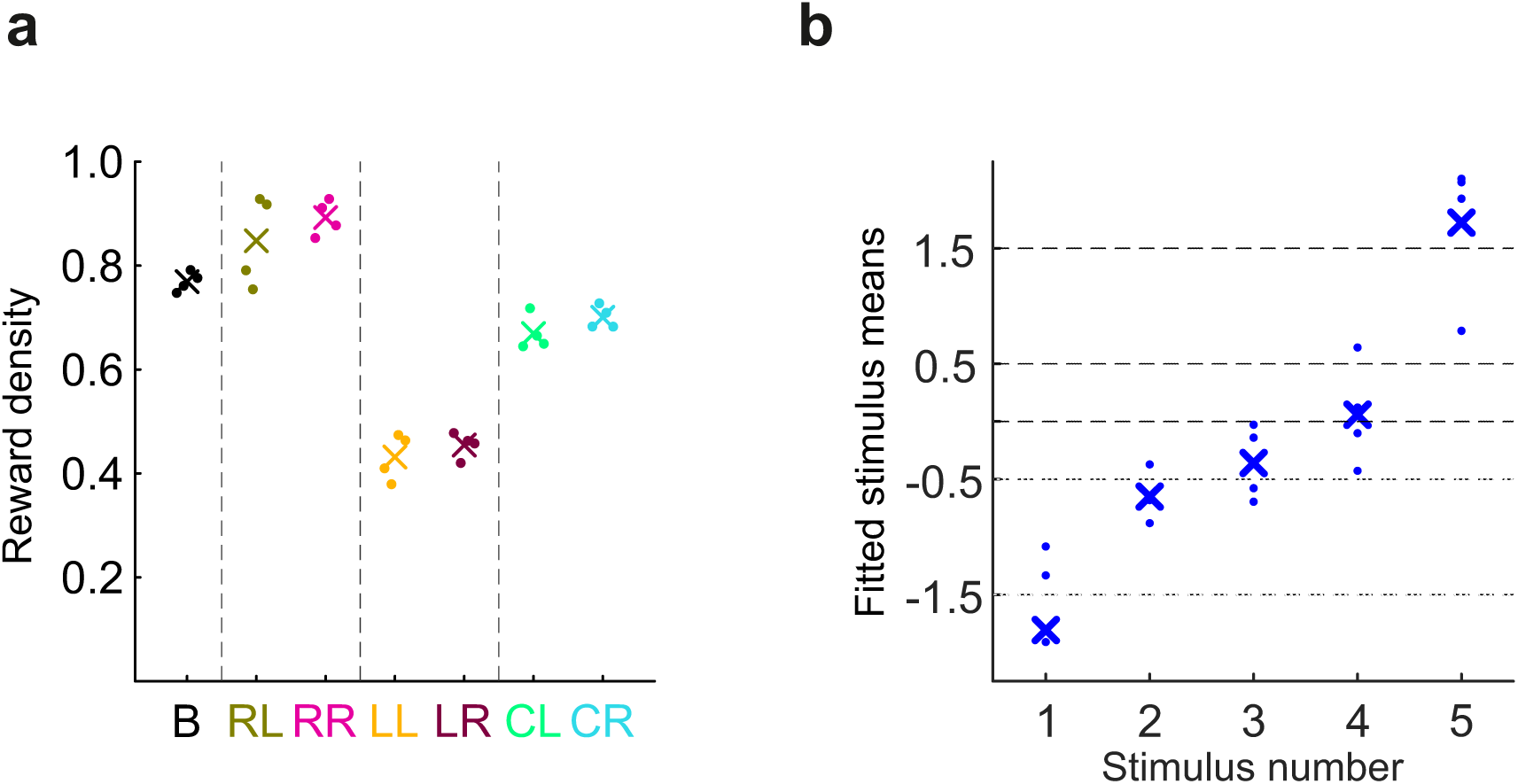
Steady-state reward densities and stimulus means. **a.** Data points represent the steady-state reward densities (i.e., average rewards per trial) computed over the last 3 sessions of each condition for each animal, whereas crosses represent means over the different animals. **b.** Individual stimulus means fitted by the OCPS model as a function of stimulus number. Individual points represent measured values for each subject across the entire experiment while crosses represent means across subjects. Dotted lines reference the intended values.

**Figure S5:**
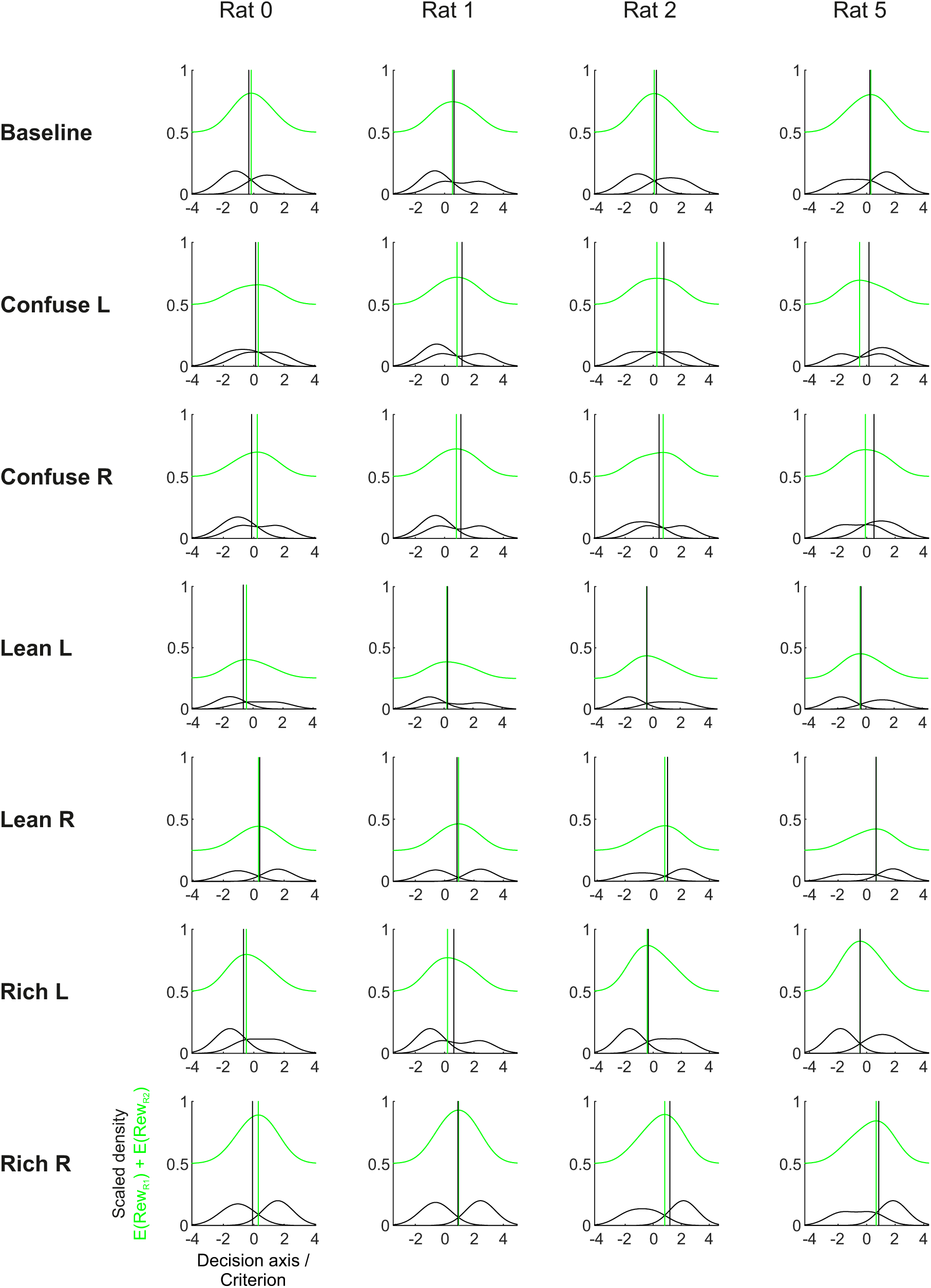
Individual steady-state vs. optimal criteria in the experimental conditions. Using the stimulus means fitted for each rat through the OCPS model, the two decision distributions (black lines) as well as the objective reward function (ORF) can be calculated. The ORF is plotted in green and its maximum, shown as a vertical green line, indicates the reward-maximizing criterion. The vertical black lines mark the animals’ steady-state criterion in the respective condition.

**Figure S6:**
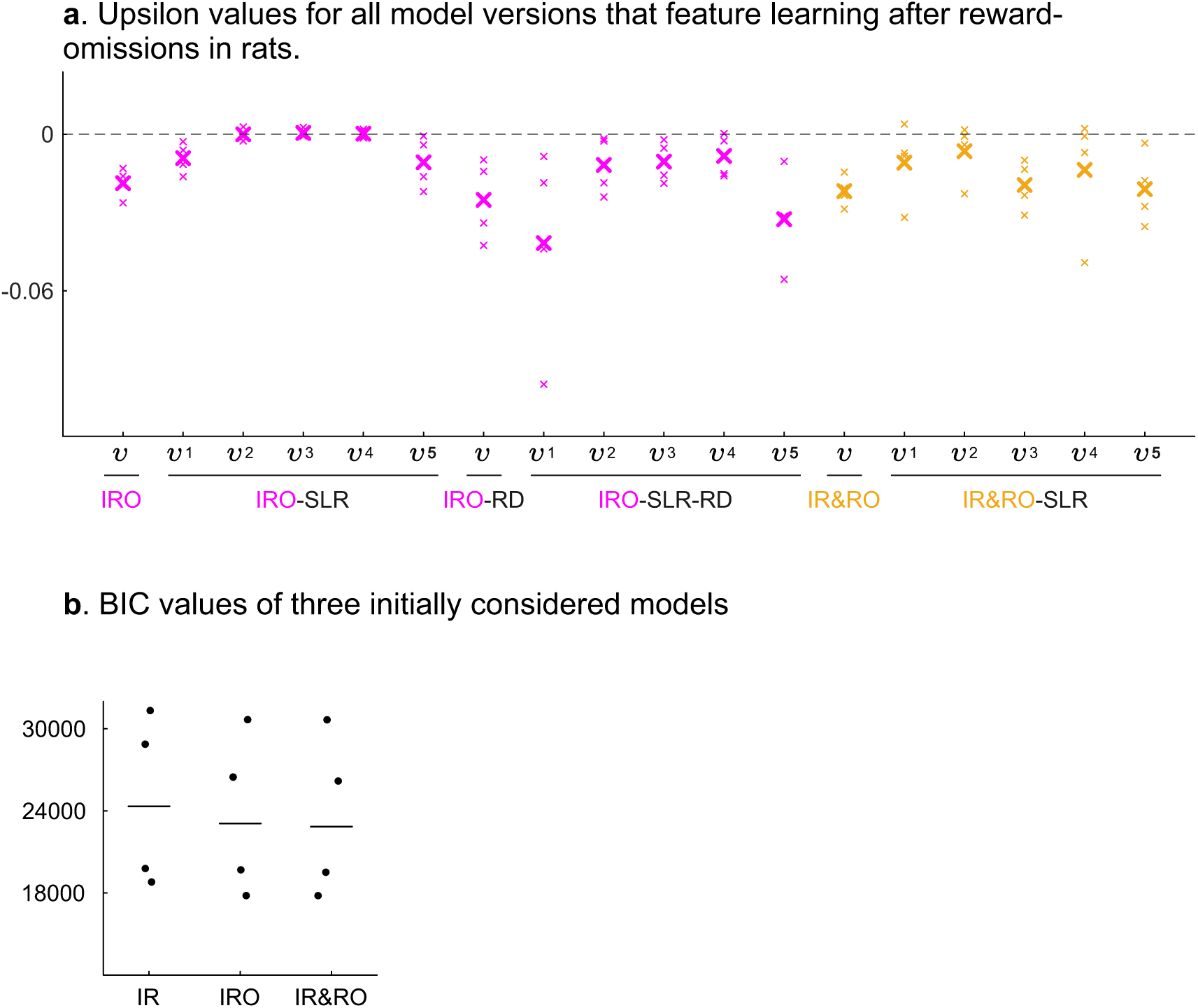
Additional figures resulting from fitting learning models to rats’ datasets. **a.** *υ* values for all model versions that feature learning after reward omissions. Both the standard IRO (pink) and IR&RO (yellow) models were extended to feature five, rather than one, stimulus-specific learning rates (SLR). Small crosses represent individual subjects’ fitted *υ* parameters, whereas thick crosses represent means over the 4 subjects. With very few exceptions, *υ* values turned out negative. **b.** Boxplots of absolute BIC values for the fits of the three initially considered models IR, IRO and IR&RO.

**Figure S7:**
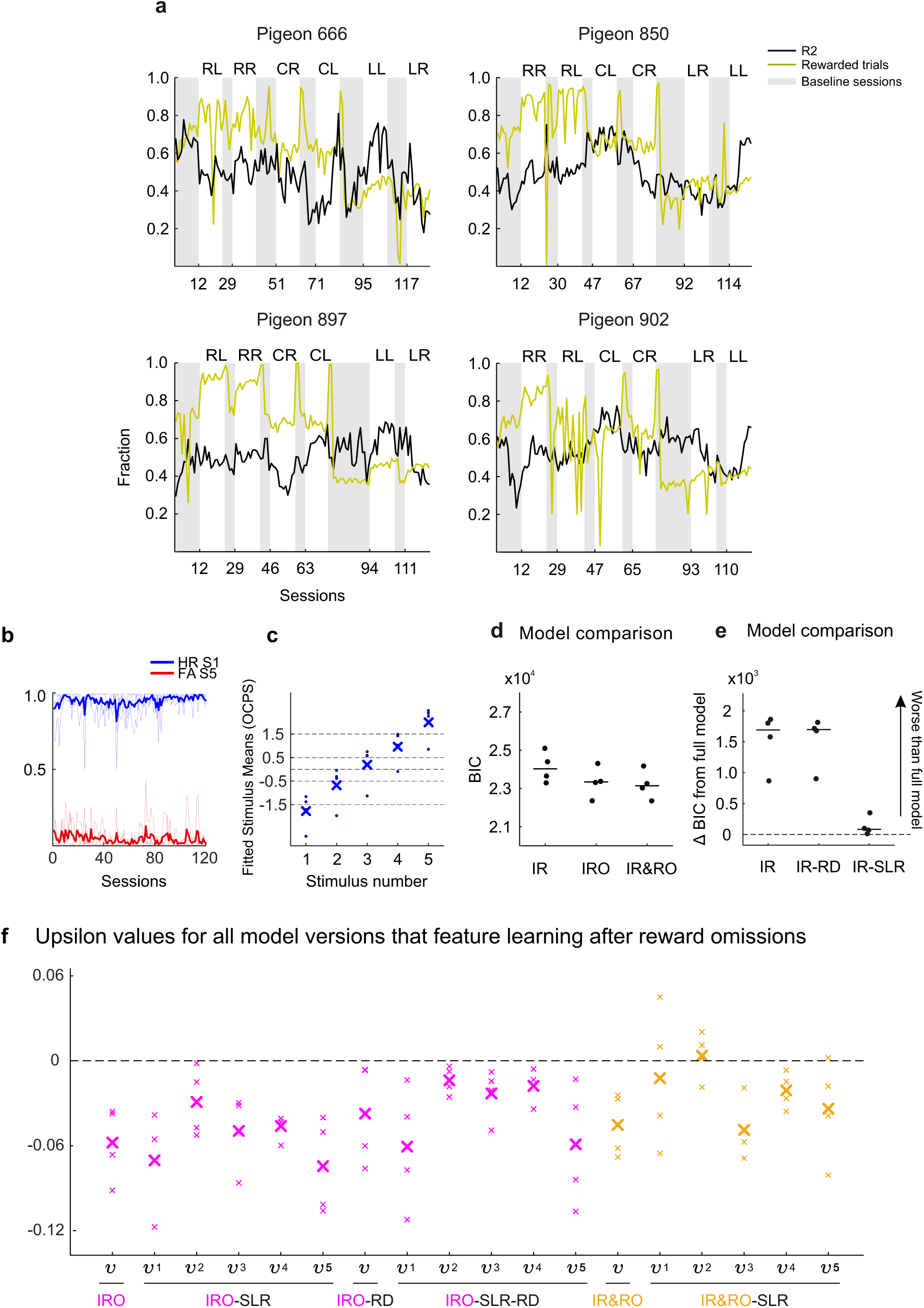
Additional results from the pigeon experiment. **a.** Session-wise fraction of R2 responses (black) and number of rewards per trial (green) for each pigeon. Gray shading indicates baseline sessions. **b.** Development of hit rate (HR, blue) for stimulus 1 and false alarm rate (FA, red) for stimulus 5 over the course of behavioral testing. Thin lines represent data from individual subjects, thick lines represent the means over the 4 subjects. **c.** Individual stimulus means fitted by the OCPS model as a function of stimulus number. Data points represent individual values while crosses represent means across subjects. Dotted lines reference the intended theoretical values. **d.** Absolute BIC values for the three initially considered models. **e.** BIC values of the competitor models after subtraction from the BIC of the full model. Unlike the full model (IR-SLR(red)-RD) used in Figure 6, which features only 2 learning rates, these BICs result from a full model version which, as in rats, indeed features 5 learning rates. In pigeons, the model with 5 learning rates systematically leads to extreme-choice behavior in Confuse conditions (see Figures S9 & S9, Pigeons 850 & 902). The full model with only two learning rates is able to fit and reproduce the data similarly and its usage leads to qualitatively identical conclusions. The three median ΔBIC values are 1691 (IR), 1697 (IR-RD), and 84.79 (IR-SLR). **f.** As in Figure S6 but for pigeons, *υ* values for all model versions that feature learning after reward omissions. Pink and yellow refer to IRO and IR&RO model versions not included in the main text.

**Figure S8:**
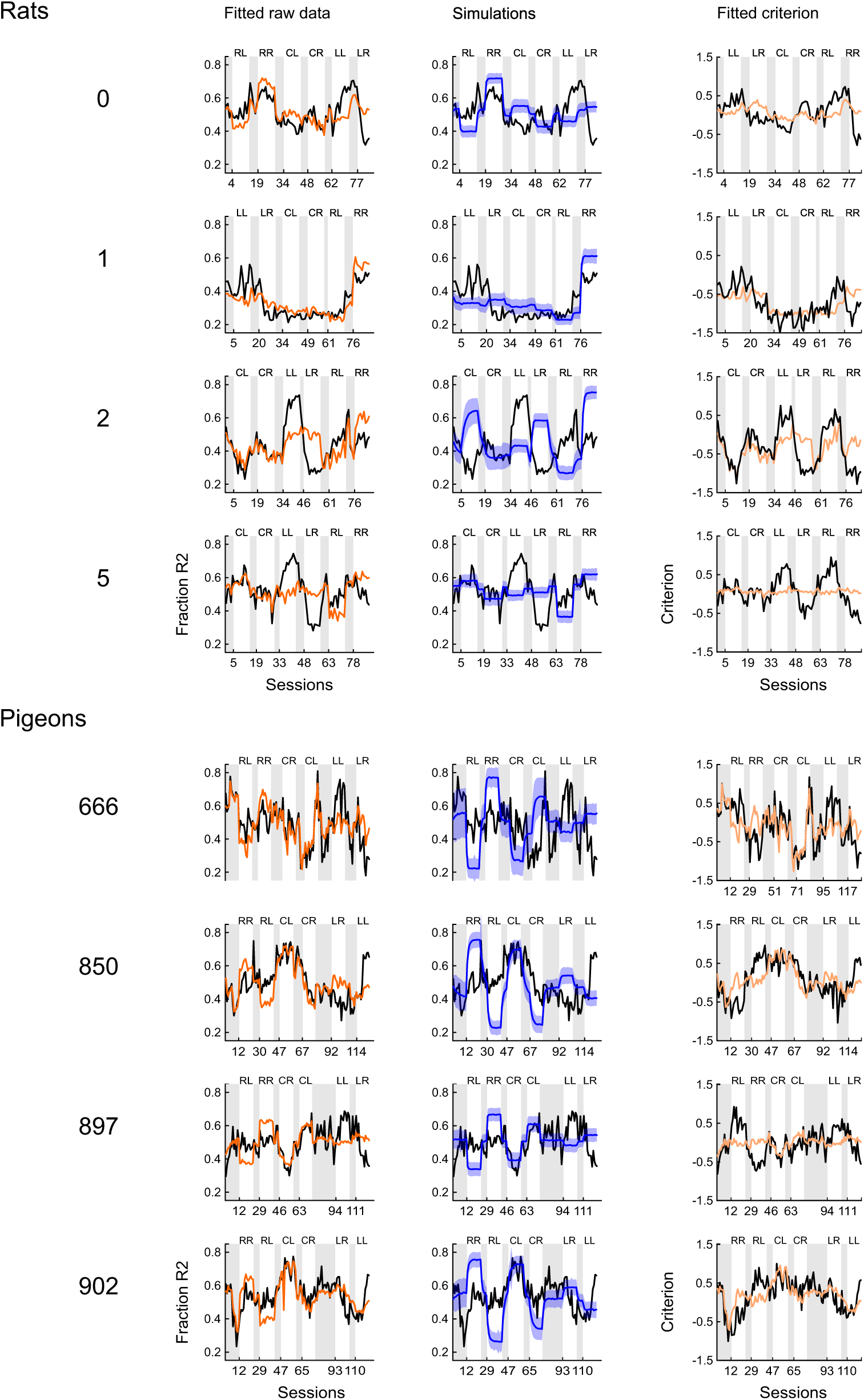
Individual fits (visualized as P(R2) and criterion) and simulations model IR for all rats and pigeons. model ***IR-SLR*** *(Integrate Rewards-Stimulus Learning Rates)*

**Figure S9:**
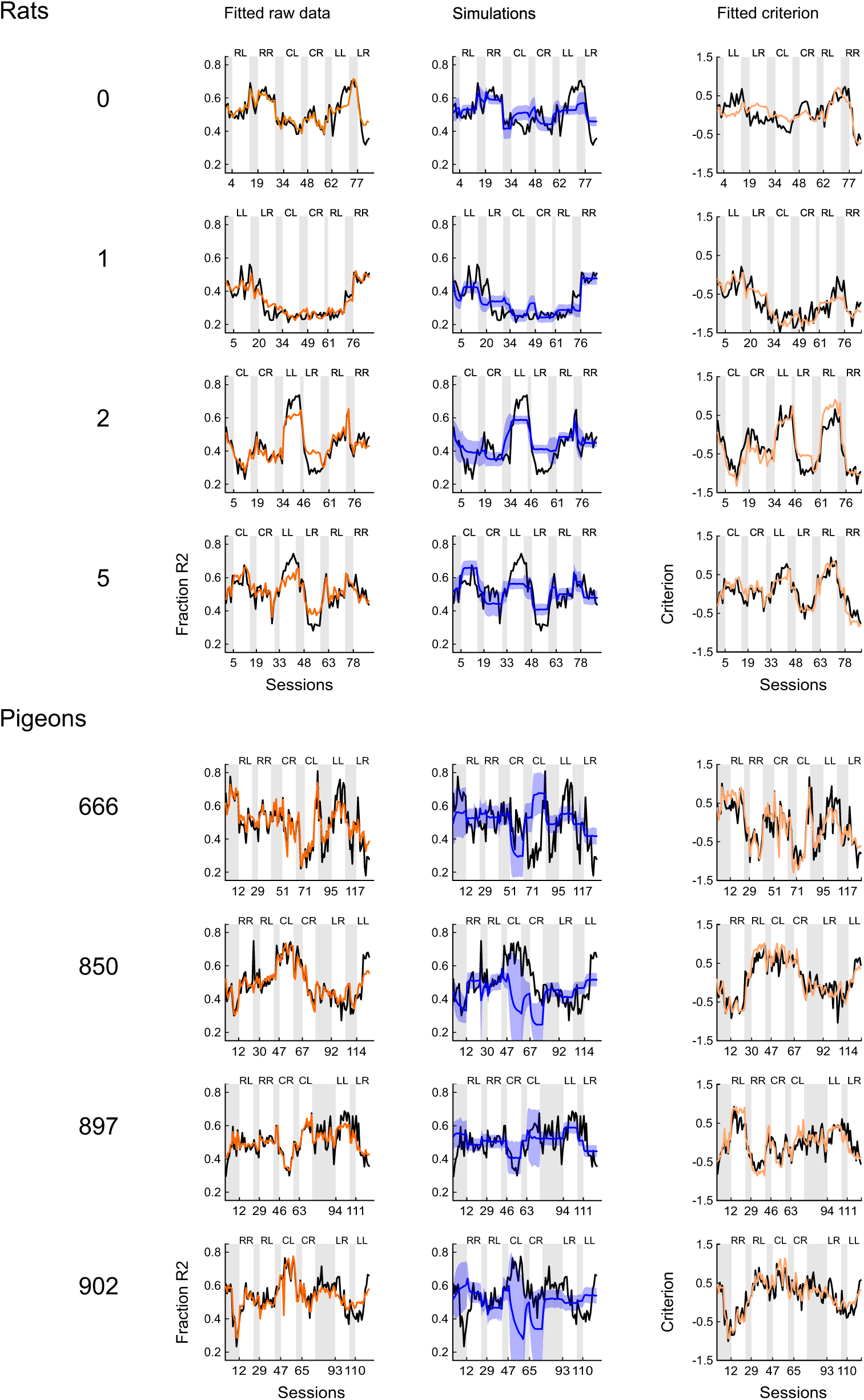
Individual fits (visualized as P(R2) and criterion) and simulations model IR-SLR for all rats and pigeons. model ***IR-RD*** *(Integrate Rewards-Relative Differences)*

**Figure S10:**
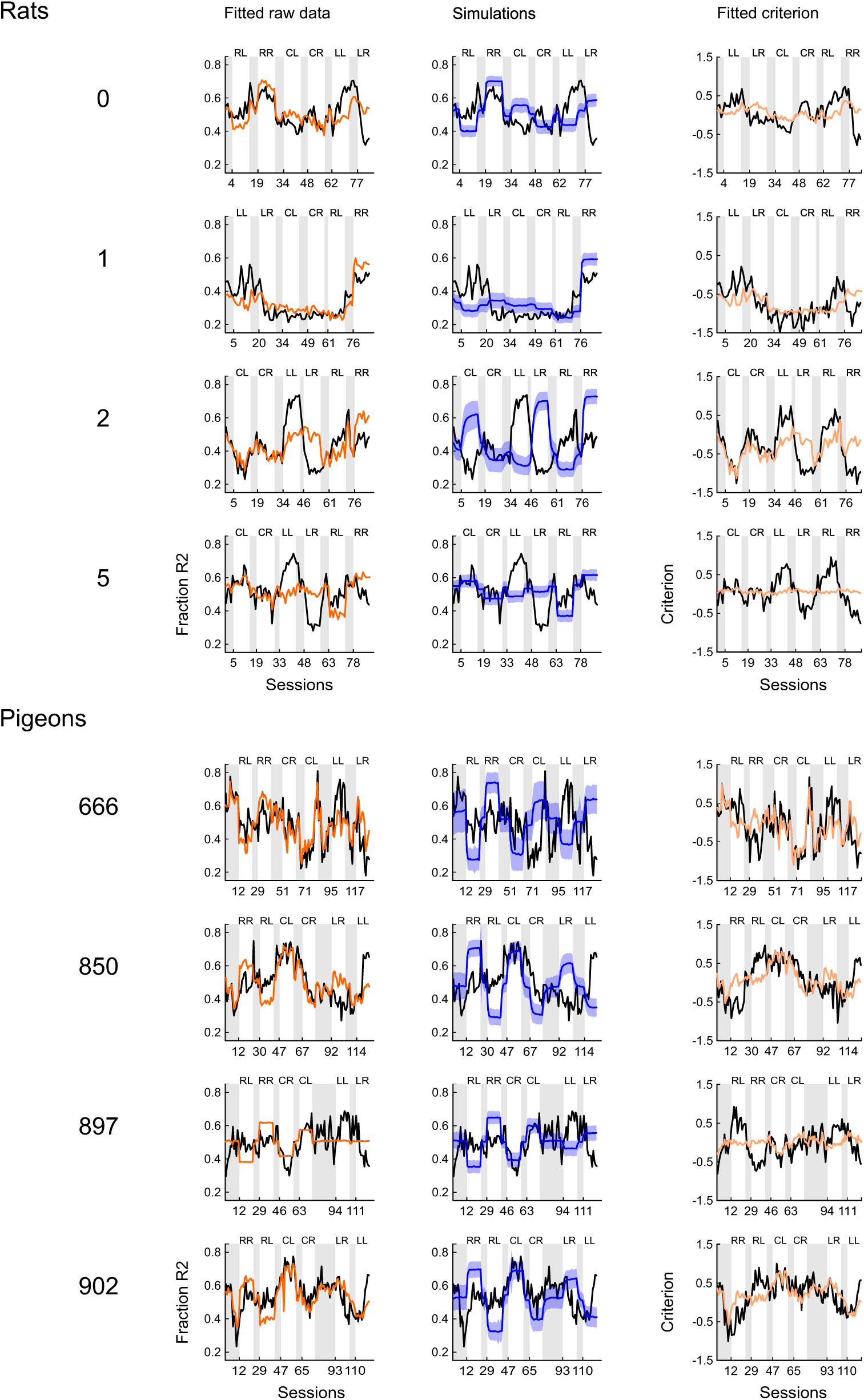
Individual fits (visualized as P(R2) and criterion) and simulations model IR-RD for all rats and pigeons. model ***IR-SLR-RD*** (*Integrate Rewards - Stimulus Learning Rates - Relative Differences)*

**Figure S11:**
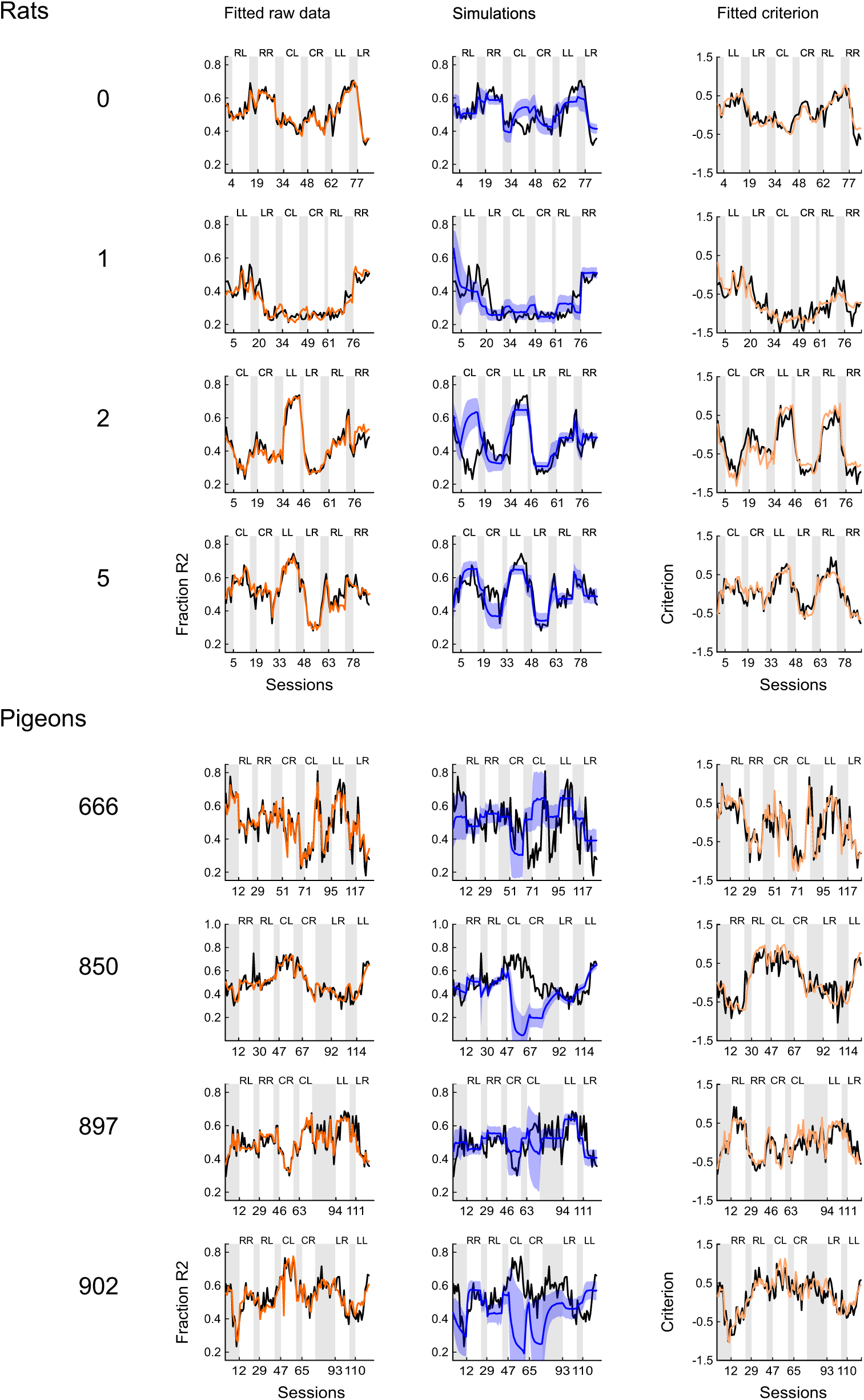
Individual fits (visualized as P(R2) and criterion) and simulations model IR-SLR-RD for all rats and pigeons. model ***IRO*** *(Integrate Reward Omissions)*

**Figure S12:**
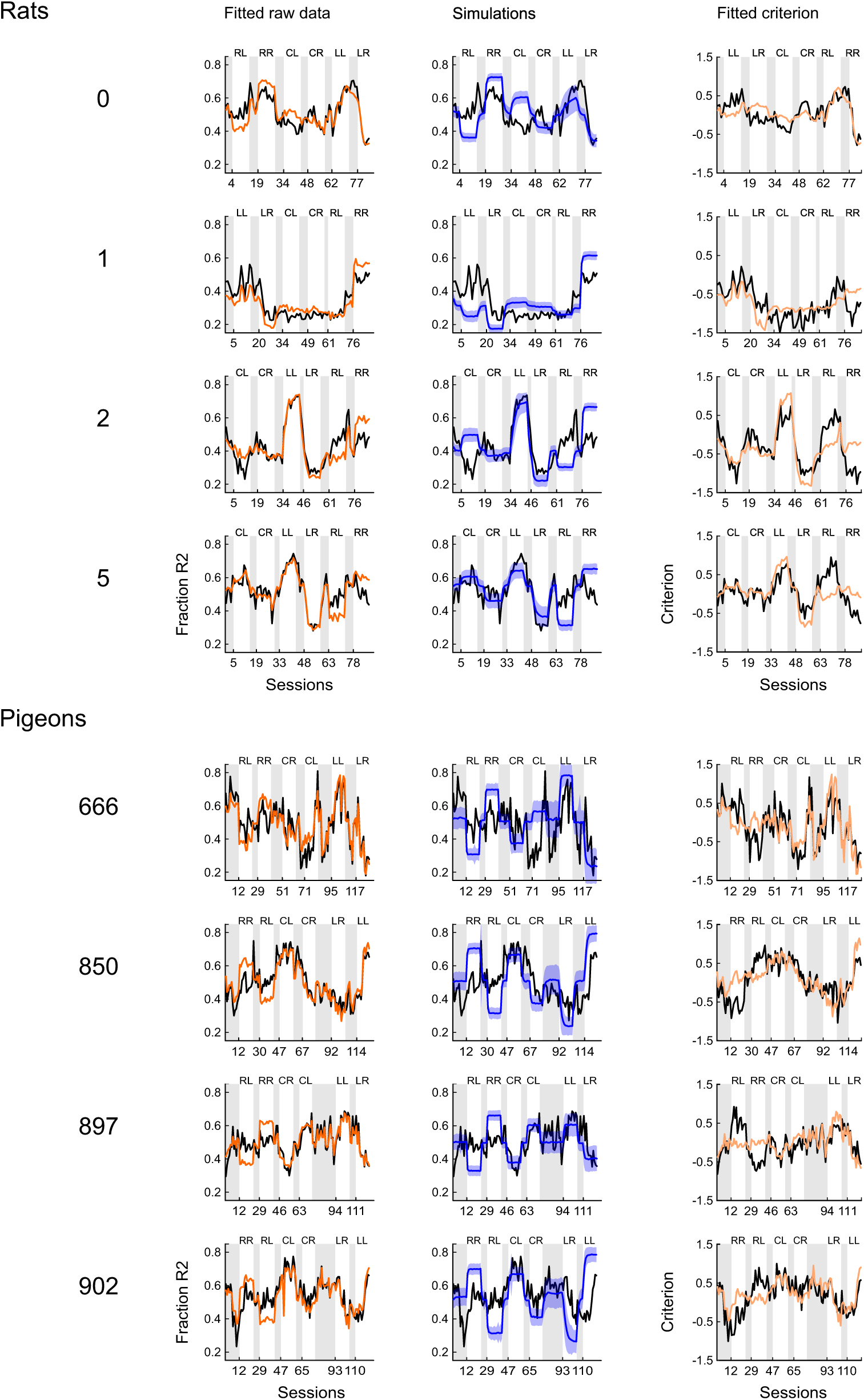
Individual fits (visualized as P(R2) and criterion) and simulations model IRO for all rats and pigeons. *model **IR&RO** (Integrate Rewards and Reward Omissions)*

**Figure S13:**
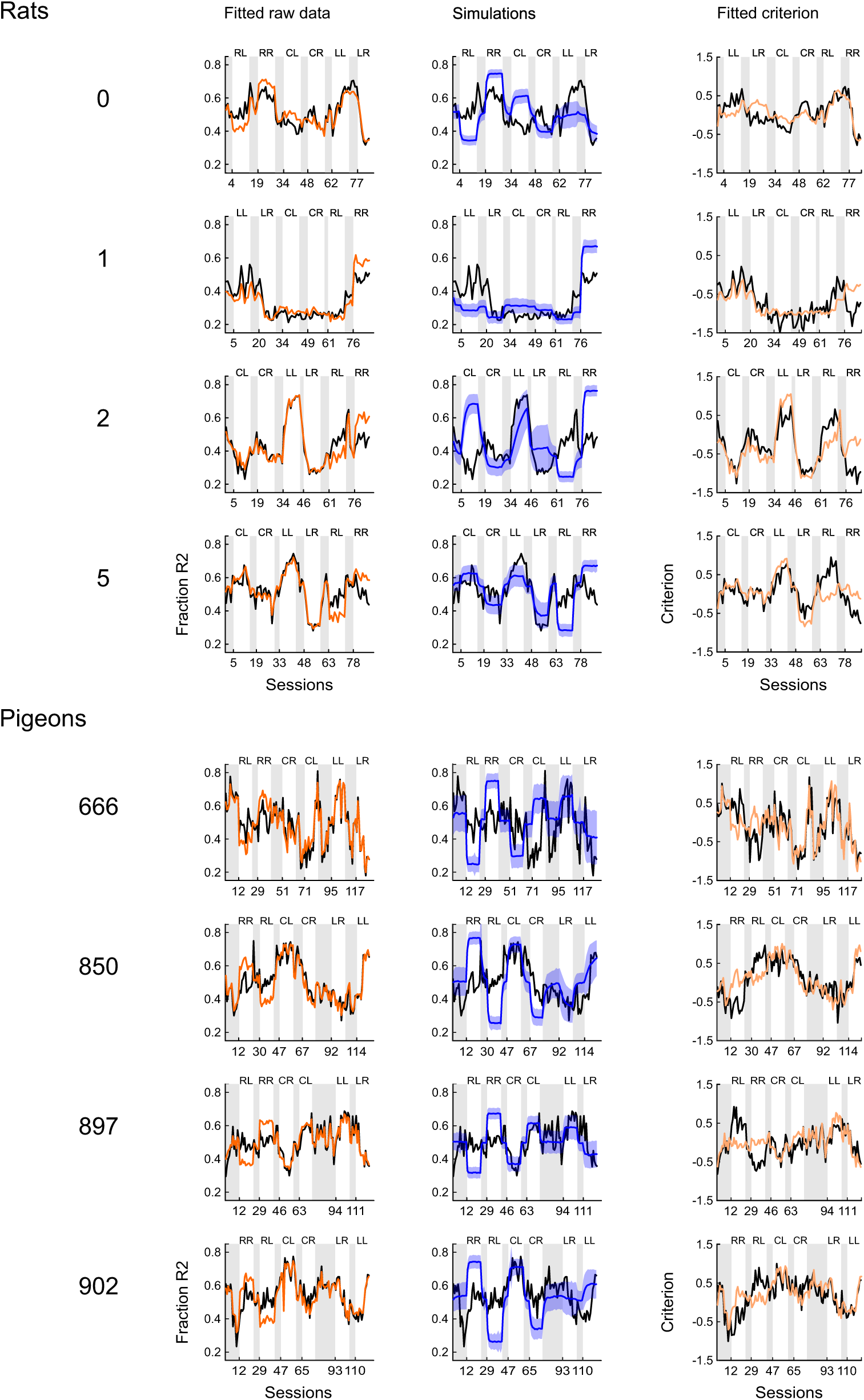
Individual fits (visualized as P(R2) and criterion) and simulations model IR&RO for all rats and pigeons. model ***IR-SLR(red)(****Integrate Rewards-Stimulus Learning Rates (reduced))* The reduced SLR version (*SLR(red)* features only 2 learning rates instead of 5 and is applied to pigeons only. See methods for details.

**Figure S14:**
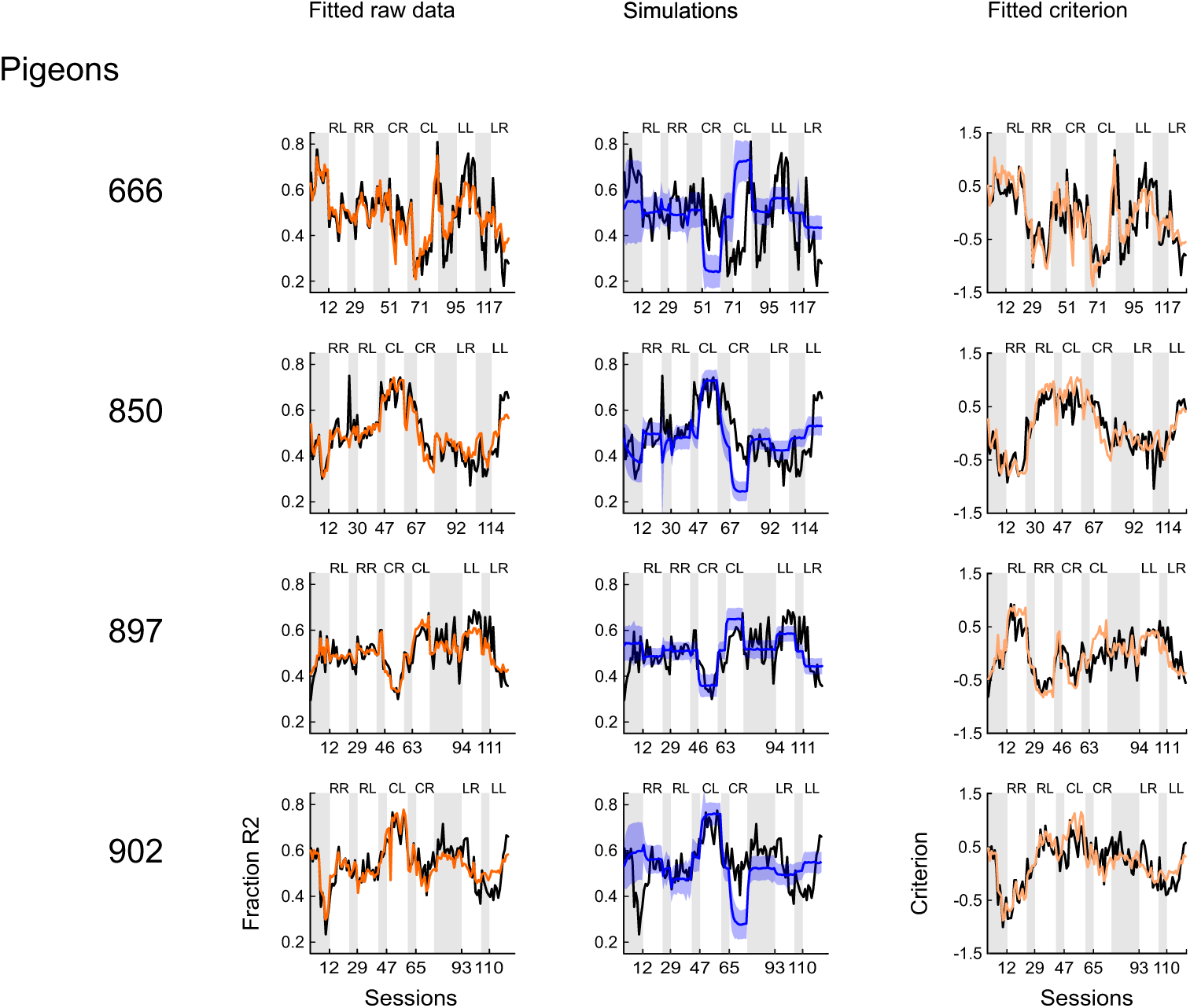
Individual fits (visualized as P(R2) and criterion) and simulations model IR-SLR(red) for all rats and pigeons. model ***IR-SLR(red)-RD*** *(Integrate Rewards - Stimulus Learning Rates (reduced) - Relative Differences)* The reduced SLR-RD version (*SLR(red)*-RD features only 2 learning rates instead of 5 and is applied to pigeons only. See methods for details.

**Figure S15:**
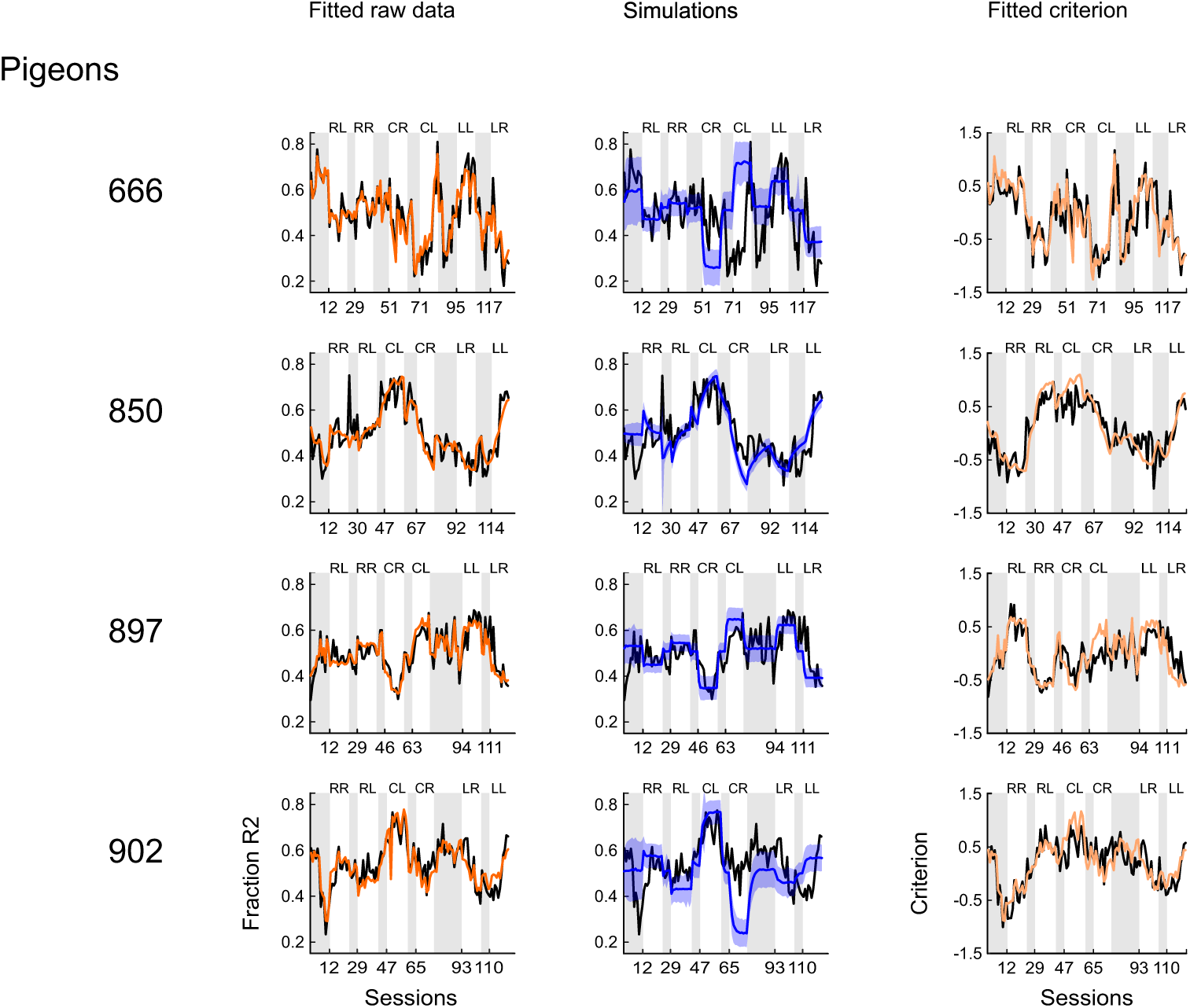
Individual fits (visualized as P(R2) and criterion) and simulations model IR-SLR(red)-RD for all rats and pigeons.

